# Organization of brainwide inputs to discrete lateral septum projection populations

**DOI:** 10.1101/2025.04.23.650257

**Authors:** Jennifer Isaac, Sonia Karkare, Hymavathy Balasubramanian, Malavika Murugan

## Abstract

The lateral septum (LS) is anatomically positioned to play a critical role in directing information from the hippocampus and cortex to downstream subcortical structures, such as the hypothalamus. In fact, early anatomical tracing studies investigated the organization of hippocampal inputs to the LS and its hypothalamic outputs to begin to understand how its structure might relate to its function. These studies also characterized the cellular anatomy of the LS and the organization of different molecular markers within it. However, relatively little is known about the organization of other, non-hypothalamic projection populations within the LS and what types of input these different projection populations receive. Here, we used retrograde tracing to determine the organization of LS projections to six different brain regions that mediate various social behaviors. We found that these projection populations occupy discrete anatomical compartments within the LS. We then used a monosynaptic rabies tracing strategy to map brainwide inputs to these six discrete LS projection populations and examine how different brain regions innervate them. We identified unique region-dependent patterns of inputs to individual LS projection populations. In particular, we observed differences in cortical, hippocampal and thalamic innervation of the six different LS projection populations, while the hypothalamic inputs were largely similar across projection populations. Thus, this study provides insight into the anatomical connectivity that may underlie the functional heterogeneity of the LS.

## Introduction

The lateral septum (LS) is a largely GABAergic subcortical structure situated ventral to the corpus callosum and medial to the lateral ventricles, which spans approximately one millimeter along the anteroposterior axis of the mouse brain, between the prefrontal cortex anteriorly and the hippocampus posteriorly^1–3^. It sends inhibitory projections to numerous subcortical structures^4^. Initially, it was suggested that the LS may act as a relay between the hippocampus and the hypothalamus, funneling distinct hippocampal inputs to distinct hypothalamic outputs to mediate a wide range of social and other motivated behaviors^4,5^. However, emerging evidence suggests a more complex role for LS^6,7^. Instead, it is proposed that the LS integrates context-related information from cortical and hippocampal inputs with information regarding internal state from subcortical inputs^8^. It is thought that distinct LS microcircuits then transform this information and transmit it to appropriate downstream structures to initiate pertinent behaviors while suppressing others^8,9^. Although the microcircuit structure of the LS has not been fully characterized, the GABAergic projection neurons of the LS are known to form local collaterals and make connections with other neurons in the LS^10^. Additionally, it has been proposed that the organization of inputs to the LS may provide insight into how the LS integrates and conveys information to its downstream targets^9^.

Early anatomical studies revealed evidence of discrete LS subregions with unique cell types^1,11^. As a result, it was first divided into dorsal, intermediate, and ventral subregions that were then further subdivided based on histological markers^12^. The inputs, especially from the hippocampus, and outputs, especially to the hypothalamus, were mapped using various tracing strategies^4,10^. Three broad classifications of cell types along the dorsoventral axis of the LS were described with dorsal LS (LSd) neurons expressing markers for somatostatin (Sst), intermediate LS (LSi) neurons expressing markers for neurotensin (Nts), and ventral LS neurons (LSv) expressing markers for estrogen receptor 1 (Esr1), among others^12,13^. Since then, studies using cell-type specific approaches have identified unique roles for these different classes of LS neurons in regulating many behaviors ranging from social approach to feeding^14–16^. However, other cell types in the LS, such as dopamine receptor 3 (Drd3) and type 2 corticotropin-releasing hormone receptor (Crhr2)-expressing neurons, have also been implicated in driving similar motivated behaviors^17,18^. Additionally, single cell sequencing and spatial transcriptomic studies have characterized novel clusters of cell types within the LS, some of which are conserved in humans^19–21^. Various LS cell types appear to have distinct developmental lineages that give rise to functional heterogeneity. For instance, perturbations of the *Nkx2.1*-lineage results in an increased exploratory phenotype^22^. Furthermore, the layered development of the LS results in a highly organized anatomical structure that likely shapes its function^23^. In fact, there is evidence of organization of kin-related social information along the dorsoventral axis of the LS^24^. Social information regarding aggressive behaviors is also organized within the LS and shaped by intraseptal circuits, as well as topographically organized hippocampal inputs^25^. Although certain social behaviors have been localized to LS subregions or cell types, it is still unclear how these subregions contribute to particular behaviors. For example, oxytocin in the LSd has been shown to both increase and attenuate anxiety associated with social fear in separate studies^26,27^. Meanwhile, oxytocin release in LSv increased aggression in female rats through inhibition of LSd^28^. One possible explanation for these mixed results is that in addition to cell-type specificity, functional differences may result from divergent input patterns onto specific subpopulations of neurons that project to distinct downstream regions^8,9^. Consistent with this explanation, a recent study found divergent roles for Nts-expressing LS neurons in driving feeding behaviors depending on where these neurons projected^29^. The Nts-expressing LS neurons that projected to the tuberal nucleus regulated hedonic feeding, while those that projected to the supramammillary nucleus regulated homeostatic feeding^29^. Interestingly, other studies have found that different populations of LSd neurons that project to the lateral hypothalamus also control feeding, with glucagon-like peptide 1 receptor (Glp1R)-expressing neurons suppressing feeding, while a certain subset of Sst-expressing neurons mediates context-dependent feeding^30,31^

A majority of studies examining LS connectivity have focused on projections to hypothalamic regions, despite evidence that the LS projects to a wide range of downstream targets that are involved in many motivated behaviors^32^. It is therefore imperative that we map input patterns to LS neurons that project beyond the hypothalamus. Long range inputs to the LS have been shown to strongly influence its involvement in a variety of behaviors^33^. One such circuit from the paraventricular hypothalamus to the LSv suppresses feeding and elicits emotional stress responses^34^. Other cortical and hippocampal projections to the LS modulate preferential investigation of a novel conspecific over a familiar conspecific^35,36^. Thus, detailed mapping of LS outputs and their monosynaptic inputs across the entire brain will help to clarify how LS circuits give rise to motivated behaviors.

Here, we chose to examine six known downstream targets of the LS that have been implicated in numerous social behaviors in distinct ways^37–40^. In particular, we describe LS projections to the basolateral amygdala (BLA) which controls social avoidance^41,42^; the bed nucleus of the stria terminalis (BNST) which is involved in the initiation of consummatory social behaviors^43^; the nucleus accumbens (NAc) which promotes social approach^44^; the periaqueductal gray (PAG) which mediates sexual and defensive behaviors^45–47^; the ventromedial hypothalamus (vmH) which regulates aggression^48^; and the ventral tegmental area (VTA) which is involved in social reward^49^. First, we sought to determine if there was topographical organization of LS outputs to these brain regions, similar to that found for hypothalamic LS projections^4^. Using a projection-specific retrograde tracing strategy, we observed organization of the six projection populations along the anatomical axes of the LS, especially along its dorsoventral axis. We then examined the organization of the brainwide inputs to each projection population using monosynaptic rabies tracing and Wholebrain mapping^50^. We found that inputs to individual LS projection populations display region-dependent specificity in innervation of different projection populations. In particular, cortical and thalamic regions show specificity in their inputs to the LS, whereas hippocampal inputs are topographically organized in a dorsoventral manner. Hypothalamic inputs are less specific, with the lateral hypothalamus innervating all six LS projection populations. We also provide evidence of reciprocal connectivity between specific hypothalamic nuclei and the LS neurons that project to those nuclei. Overall, the organization of various projection populations within the LS and the differing patterns of inputs they receive likely contributes to the functional heterogeneity of the LS.

## Results

### Anatomical organization of discrete projection populations within the lateral septum

In order to determine the anatomical organization of projection populations within the LS, we implemented a retrograde viral tracing strategy in which we injected a retrogradely transported Cre virus labeled with an mCherry fluorophore into one of six downstream brain regions and a Cre-dependent green fluorescent protein (GFP) into the LS (Figure 1A). Once expression of the retro Cre was confirmed in the downstream region of interest (Supplementary Figure 1), the GFP-labeled neurons in the LS (Figure 1B; Supplementary Figure 2) were quantified across the anteroposterior (A-P) axis of the LS (Supplementary Figure 3A). The quantification of GFP-labeled LS neurons was performed by registering approximately 50 μm tissue sections containing the LS from 1.3 mm anterior to bregma to 0.3 mm posterior to bregma to the corresponding section from the Allen Brain Atlas using Wholebrain software (Figure 1C, left panel; Supplementary Figure 3B). GFP-labeled LS neurons on the side ipsilateral to the viral injection were then semi-automatically detected and manually confirmed (Figure 1C, right panel). All other neurons, including those in the medial septum and contralateral LS, were excluded from analyses. This enabled us to create a detailed map of individual LS projection neurons across three animals per projection population (Figure 1E). We found that varying numbers of LS neurons project to the different downstream brain regions (Figure 1D). In particular, LS neurons that project to the NAc are the most numerous (LS-NAc average number of neurons: 6171.33 ± 458.02), followed by neurons that project to the vmH (LS-vmH average number of neurons: 1488 ± 453.10) and BNST (LS-BNST average number of neurons: 966 ± 167.48), with the BLA (LS-BLA average number of neurons: 589.67 ± 275.12), VTA (LS-VTA average number of neurons: 439.67 ± 148.95) and PAG (LS-PAG average number of neurons: 97.67 ± 30.24) receiving fewer inputs from the LS (one-way ANOVA with post-hoc t tests, p = 9.70*10^-11^, see Supplementary Table 1). The LS has historically been divided into subregions: rostral LS (LSr), caudal LS (LSc), ventral LS (LSv), septohippocampal nucleus (SH) and septofimbrial nucleus (SF)^1,12^. Our LS neuron mapping strategy allows us to quantify the proportion of projection neurons in each of these divisions. We found that the LS-NAc projection consists of more LSc neurons than the LS-BNST, LS-vmH or LS-VTA populations, and the LSr was the greatest input to the LS-BLA, LS-BNST, LS-vmH and LS-VTA projection populations. (Supplementary Figure 3C, two-way ANOVA with projection population and LS subregion as factors, interaction: p = 0.001, projection population: p = 1, LS subregion: p = 9.97*10^-25^ with post-hoc t tests, see Supplementary Table 1). While these subdivisions provide useful anatomical boundaries, there is evidence to suggest that each of these subregions can be further divided based on histological markers and connectivity mapping^12,13^. Thus, for our subsequent analyses, we collapsed data across all LS subdivisions and quantified the distribution of LS projection neurons based on their anatomical coordinates.

**Figure 1.**
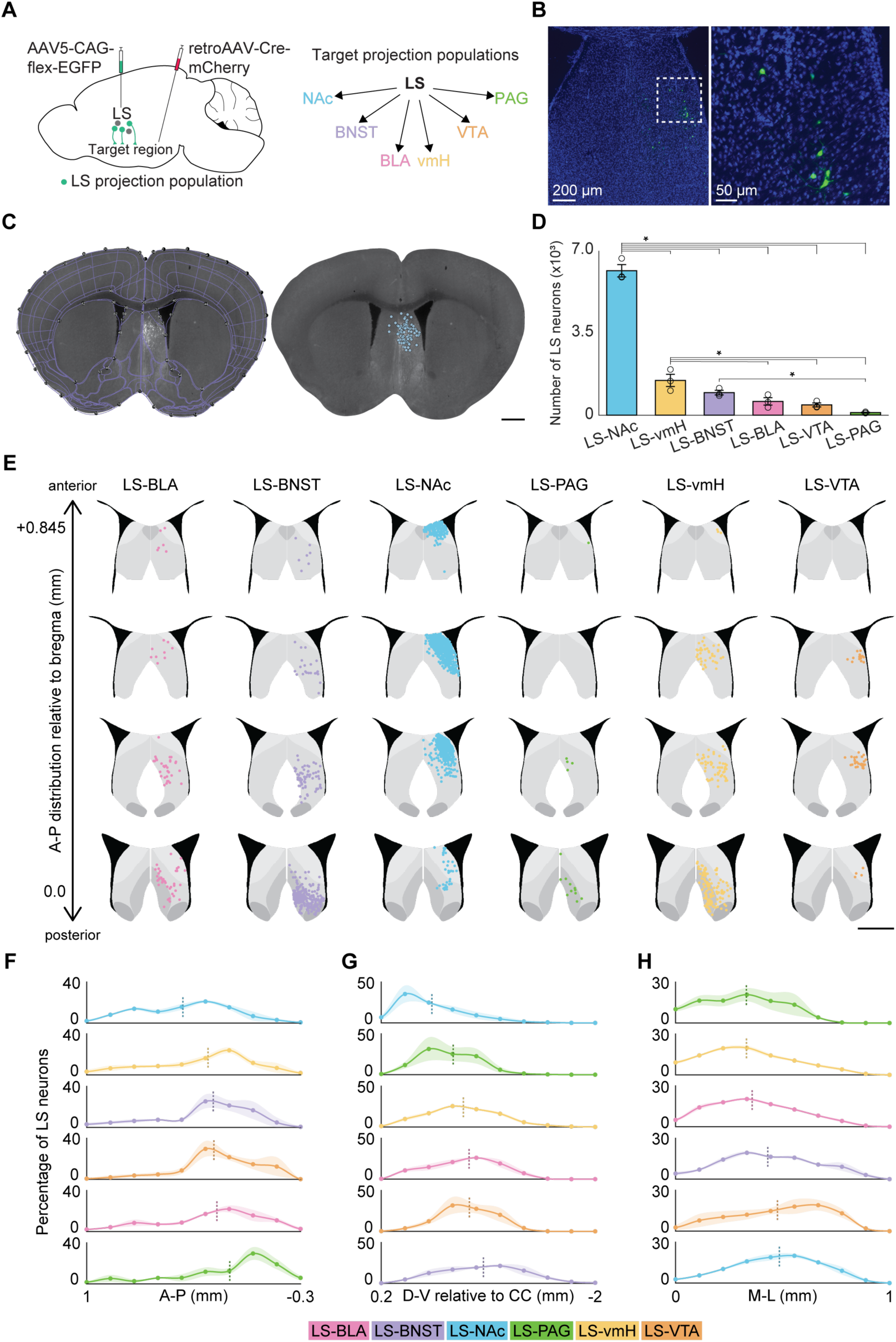
Organization of projection population neurons along various anatomical axes of the lateral septum. A) Schematic of the retrograde viral tracing strategy used to label LS neurons that project to a target brain region with GFP and neurons within the target brain region with mCherry (left panel). The six target brain regions (right panel) are the basolateral amygdala (BLA), bed nucleus of the stria terminalis (BNST), nucleus accumbens (NAc), periaqueductal gray (PAG), ventromedial hypothalamus (vmH), and ventral tegmental area (VTA). B) Example histology from a representative animal showing LS projection population neurons retrogradely labeled with GFP at 10x (left panel) and 20x (right panel) magnification. GFP shown in green and DAPI shown in blue. Scale bar represents 200 μm (left panel, 10x) and 50 μm ( right panel, 20x). C) Schematic of the Wholebrain registration process. Gray-scaled coronal sections of the LS are registered to match corresponding sections from the Allen Brain Atlas (left panel). Neurons labeled with GFP are identified using a filter for soma size and brightness. They are then assigned to their region identity based on registration to the atlas (right panel). Scale bar represents 1 mm. D) Cell counts of LS projection populations show that the LS-NAc is the largest of the LS projection populations, followed by the LS-vmH and LS-BNST projections. One-way ANOVA (p = 9.70*10^-11^) with post-hoc t tests (LS-NAc to LS-vmH: 2.57*10^-9^, LS-NAc to LS-BNST: 6.54*10^-10^, LS-NAc to LS-BLA: 2.49*10^-10^, LS-NAc to LS-VTA: 1.69*10^-10^, LS-NAc to LS-PAG: 7.03*10^-11^, LS-vmH to LS-BLA: 0.030, LS-vmH to LS-VTA: 0.011, LS-vmH to LS-PAG: 0.001, LS-BNST to LS-PAG: 0.037). E) Example sections of identified LS neurons projecting to each of the target brain regions across the A-P axis of the LS from anterior (top) to posterior (bottom). Compartments of the lateral septal complex (LSX) are colored in varying shades of gray. Each column consists of one representative animal from each projection population, and neurons are color coded by projection population (LS-BLA: pink, LS-BNST: purple, LS-NAc: blue, LS-PAG: green, LS-vmH: yellow, LS-VTA: orange). Scale bar represents 1 mm. F) Distributions of LS projection populations across the A-P axis of the LS. Projections are organized from anterior to posterior based on their median A-P coordinate, denoted by the dashed line (LS-NAc: 0.415, LS-vmH: 0.262, LS-BNST: 0.231, LS-VTA: 0.227, LS-BLA: 0.208, LS-PAG: 0.103), with the LS-NAc being the most anterior population and the LS-PAG being the most posterior population. Two-way ANOVA with projection population and bin as factors (interaction: p = 1.36*10^-3^, projection population: p = 1, bin: p = 3.73*10^-23^) with post-hoc t tests (see Supplementary Table 1). G) Distributions of LS projection populations across the D-V axis of the LS relative to the corpus callosum (CC). Projection populations are organized from dorsal to ventral based on their median D-V coordinate relative to the CC, denoted by the dashed line (LS-NAc: −0.320, LS-PAG: −0.539, LS-vmH: −0.645, LS-BLA: −0.704, LS-VTA: −0.707, LS-BNST: −0.851), with the LS-NAc being the most dorsal population and the LS-BNST being the most ventral population. Two-way ANOVA with projection population and bin as factors (interaction: p = 1.28*10^-5^, projection population: p = 1, bin: p = 6.72*10^-28^) with post-hoc t tests (see Supplementary Table 1). H) Distributions of LS projection populations across the M-L axis of the LS. Projections are organized from medial to lateral based on their median M-L coordinate, denoted by the dashed line (LS-PAG: 0.331, LS-vmH: 0.332, LS-BLA: 0.358, LS-BNST: 0.431, LS-VTA: 0.476, LS-NAc: 0.485), with the LS-PAG and LS-vmH being the most medial populations and the LS-VTA and LS-NAc being the most lateral populations. Two-way ANOVA with projection population and bin as factors (interaction: p = 1.96*10^-5^, projection population: p = 0.999, bin: p = 6.33*10^-34^) with post-hoc t tests (see Supplementary Table 1). Values from individual animals indicated by unfilled circles. Error bars and shaded error regions indicate ± SEM. *p < 0.05. Coordinates are relative to bregma unless otherwise noted. Representative histology shown in B is from the LS-VTA projection population and representative histology shown in C is from the LS-vmH projection population. N = 3 animals per projection population.

Using the maps we generated of individual LS projection populations, we examined the organization of these projection populations along the various anatomical axes of the LS. We determined the distribution of the proportion of LS neurons of each projection population across the A-P (Figure 1F), dorsoventral (D-V, Figure 1G), and mediolateral (M-L, Figure 1H) axes. The D-V position is reported as distance relative to the corpus callosum to account for the changing depth of the LS along the A-P axis. The average median value of each distribution can be organized in an anterior to posterior, dorsal to ventral and medial to lateral manner. Along the A-P axis, the median A-P coordinates (relative to bregma in mm) of each projection population are distributed from anterior to posterior as follows: LS-NAc: 0.415, LS-vmH: 0.262, LS-BNST: 0.231, LS-VTA: 0.227, LS-BLA: 0.208, LS-PAG: 0.103. Along the D-V axis, the median D-V coordinates (relative to the corpus callosum in mm) of each projection population are distributed from dorsal to ventral as follows: LS-NAc: −0.320, LS-PAG: −0.539, LS-vmH: −0.645, LS-BLA: −0.704, LS-VTA: −0.707, LS-BNST: −0.851. Along the M-L axis, the median M-L coordinates (relative to bregma in mm) of each projection population are distributed from medial to lateral as follows: LS-PAG: 0.331, LS-vmH: 0.332, LS-BLA: 0.358, LS-BNST: 0.431, LS-VTA: 0.476, LS-NAc: 0.485.

The LS has a complex anatomical structure that varies along its A-P axis (Figure 1F; Supplementary Figure 2; Supplementary Figure 3A). As a result, we sought to determine how the individual projection populations varied in their anatomical location at specific A-P coordinates (0.845 mm, 0.545 mm and 0.02 mm anterior to bregma) that span the A-P axis of the LS. At each of these A-P coordinates, we created density contour plots which show the distribution of LS neurons collapsed across three animals per projection population (Figure 2A, C and E). At the most anterior A-P coordinate (0.845 mm), comparison of the 90% contours revealed significant differences in the distribution of the LS-BNST population from all other LS populations (Figure 2A, B; pairwise KS tests, LS-BNST to LS-BLA: p = 6.16*10^-4^, LS-BNST to LS-NAc: p = 8.94*10^-5^, LS-BNST to LS-PAG: p = 0.00107, LS-BNST to LS-vmH: p = 2.48*10^-4^, LS-BNST to LS-VTA: p = 3.80*10^-4^, see Supplementary Table 1). At the intermediate coordinate (0.545 mm), all populations are differentially distributed relative to other populations (Figure 2C, D; pairwise KS tests, see Supplementary Table 1). At the more posterior A-P coordinate (0.02 mm), pairwise KS tests can discriminate between the 90% contours of all projection populations except for the LS-BLA and LS-vmH projections which are more anatomically overlapping in posterior LS (Figure 2E, F; pairwise KS tests, see Supplementary Table 1). The anatomical organization of these projection populations is further demonstrated by plotting the D-V and M-L coordinate of the centroid of the density contours (Figure 2B, D and F). At the anterior A-P coordinate (Figure 2B), centroids are clustered together along the D-V axis with the exception of the LS-BNST (centroid D-V coordinate - LS-BLA: −0.08, LS-BNST: −0.37, LS-NAc: −0.11, LS-PAG: −0.08, LS-vmH: −0.17, LS-VTA: −0.18). At the intermediate A-P coordinate (Figure 2D), the centroids are organized along the D-V axis (centroid D-V coordinate - LS-BLA: −0.22, LS-BNST: −0.78, LS-NAc: −0.11, LS-PAG: −0.27, LS-vmH: −0.51, LS-VTA: −0.41). At the posterior A-P coordinate (Figure 2F), centroids are still relatively organized along the D-V axis (centroid D-V coordinate - LS-BLA: −0.84, LS-BNST: −1.2, LS-NAc: −0.70, LS-PAG: −0.27, LS-vmH: −0.83, LS-VTA: −1.02) and display increased organization along the M-L axis (centroid M-L coordinate - LS-BLA: 0.22, LS-BNST: 0.31, LS-NAc: 0.71, LS-PAG: 0.51, LS-vmH: 0.27, LS-VTA: 0.56). The anatomical organization of these projection populations appears qualitatively similar to the spatial mapping of various cell markers^12,13,16,19^, raising the possibility that LS projection populations may correspond to specific LS cell types.

**Figure 2.**
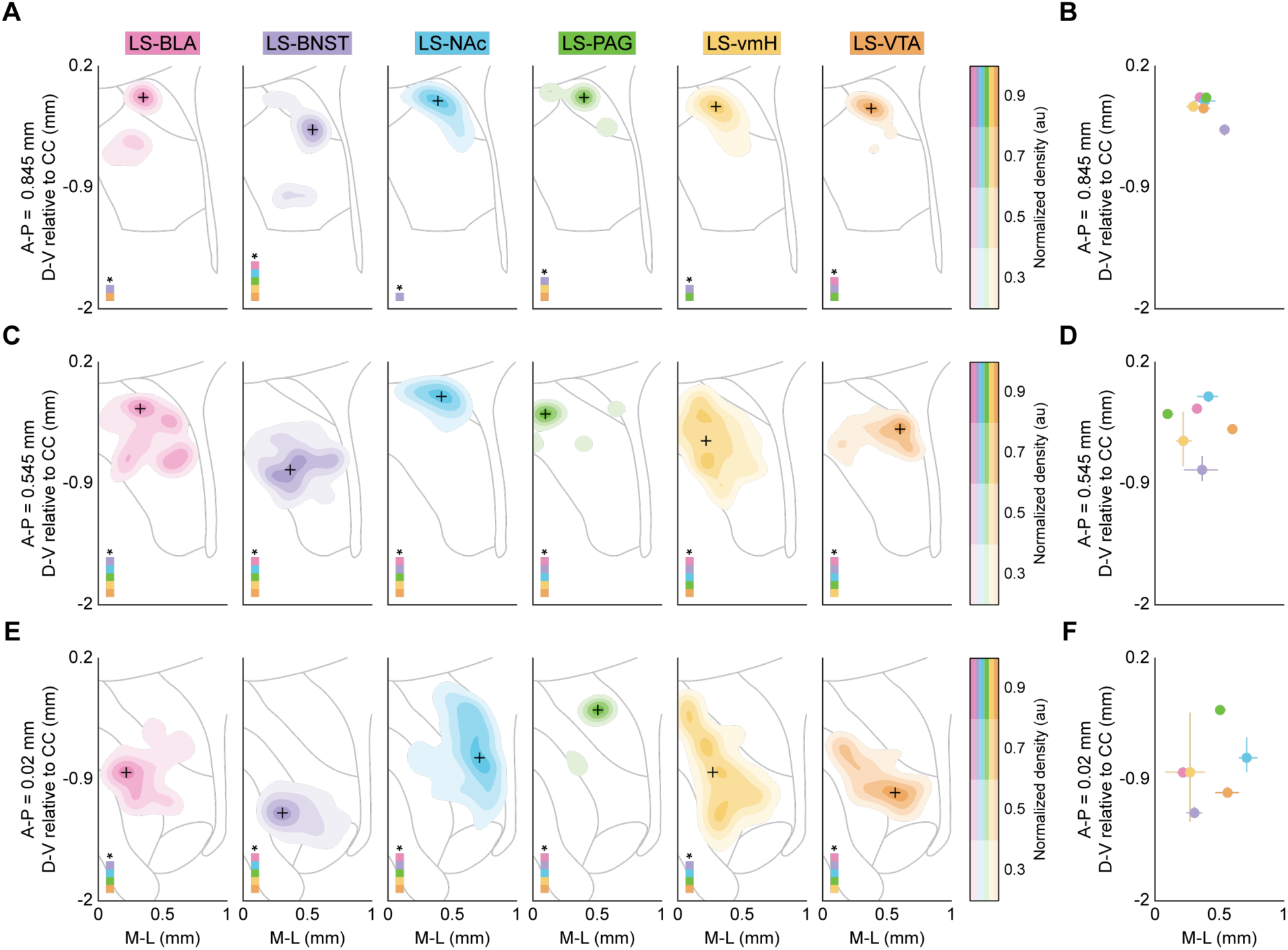
Projection populations occupy discrete regions along the D-V and M-L axes of the LS. A) Density contour plots depicting the location of projection neurons within the LS across all animals from each projection population at an A-P coordinate of 0.845 mm anterior to bregma. The gray outline corresponds to the LS, including the SH, LSc and LSr, at an A-P of 0.845 mm with the partial outline of the ventricle. At this A-P, comparison of the 90% contour distributions demonstrates that the LS-BNST population is the only distribution to significantly differ from all other LS projection populations, while the LS-VTA differs from the LS-BLA and LS-PAG and the LS-PAG differs from the LS-vmH (see Supplementary Table 1). B) Centroids of 90% contours plotted with the horizontal and vertical spread of the 90% contour at an A-P of 0.845 mm. At this A-P, the LS-BNST population is located more ventrally and laterally than all other LS projection populations, which are clustered in a more medial and dorsal location. C) Density contour plots depicting the location of projection neurons within the LS across all animals from each projection population at an A-P coordinate of 0.545 mm anterior to bregma. The gray outline corresponds to the LS, including the LSc and LSr, at an A-P of 0.545 mm with the partial outline of the ventricle. At this A-P, comparison of the 90% contour distributions using a pairwise KS test demonstrates that all LS projection populations are significantly different from each other (see Supplementary Table 1). D) Centroids of 90% contours plotted with the horizontal and vertical spread of the 90% contour at an A-P of 0.545 mm. At this A-P, LS projection populations are separable based on their centroid and spread. All populations except the LS-BNST and LS-vmH projections have little spread, with the LS-NAc, LS-BLA, and LS-PAG populations being most dorsally located. The LS-VTA and LS-vmH populations have intermediately positioned centroids, while the LS-BNST population is the most ventral. E) Density contour plots depicting the location of projection neurons within the LS across all animals from each projection population at an A-P coordinate of 0.02 mm anterior to bregma. The gray outline corresponds to the LS, including the LSc, LSr, LSv and SF at an A-P of 0.545 mm with the partial outline of the ventricle and fornix. At this A-P, comparison of the 90% contour distributions demonstrates that all LS projection populations except for the LS-BLA and LS-vmH populations are separable based on a pairwise KS test (see Supplementary Table 1). F) Centroids of 90% contours plotted with the horizontal and vertical spread of the 90% contour at an A-P of 0.02 mm. At this A-P, most LS projection populations are separable based on their centroid and spread, with the centroids of the LS-PAG and LS-NAc populations being more dorsally located, while the LS-BNST population remains more ventrally positioned. The LS-BLA, LS-vmH, and LS-VTA populations are more intermediate, though the LS-vmH population has a large D-V spread. Each concentric layer of the contour plots corresponds to a normalized density of 30%, 50%, 70% and 90% (from lightest to darkest) in the color corresponding to individual projection populations. The colored squares in the bottom left of each panel indicate significant differences between the corresponding projection populations based on a pairwise KS test (see Supplementary Table 1). *p < 0.05. N = 3 animals per projection population.

### Specific LS projection populations receive different proportions of inputs from major brain regions

There is evidence that the organization of inputs into the LS could influence how information is integrated by LS subpopulations^8,9^. After we identified discrete projection populations organized anatomically within the LS, we next characterized the monosynaptic inputs to these projection populations using a modified rabies tracing strategy. We first isolated projection populations by injecting a retrogradely transported Cre virus into one of the six downstream brain regions along with a Cre-dependent helper virus labeled with GFP into the LS (Figure 3A, left panel). After three weeks, we injected a modified rabies virus labeled with mCherry into the LS (Figure 3A, right panel) to identify the brainwide monosynaptic inputs to individual projection populations. We then registered approximately 50 μm sections to the corresponding section in the Allen Brain Atlas across the entire A-P axis (2.545 mm anterior to bregma to 4.955 mm posterior to bregma) using Wholebrain software (Figure 3C, Supplementary Figure 4) of three animals per projection population. We confirmed the presence of starter cells, those co-labeled with GFP (Cre-dependent helper virus) and mCherry (modified rabies virus), in the LS (Figure 3B). After tracking all monosynaptic input neurons across the entire brain, we determined the distribution of the proportion of neurons along the A-P axis (Figure 3D). These distributions were largely similar across projection populations (two-way ANOVA with projection population and bin as factors, interaction: p = 0.88, projection: p = 1, bin: p = 3.43*10^-53^ with post-hoc t tests, see Supplementary Table 1). We then quantified the proportion of inputs from major brain regions as defined by the Allen Brain Atlas (Figure 3E, see ‘Quantification of brainwide inputs’ under *Methods* section). We found that LS-BLA neurons receive the greatest input from the hippocampal formation (35.04% ± 11.15%), and similar proportions of inputs from the isocortex (20.69% ± 7.80%) and the striatum (16.58% ± 13.89%). LS-BNST neurons receive similar proportions of inputs from the hippocampal formation (25.89% ± 11.77%) and isocortex (26.66% ± 11.20%) and lower proportions of inputs from the midbrain (4.89% ± 2.95%) and pallidum (3.91% ± 0.94%). LS-NAc neurons receive the greatest proportion of inputs from the hippocampal formation (29.86% ± 3.31%) and isocortex (29.79% ± 1.41%). LS-PAG neurons receive a significantly larger proportion of their inputs from the isocortex (45.39% ± 8.35%) relative to all other regions. LS-vmH neurons receive the greatest proportion of inputs from the hippocampal formation (35.02% ± 3.12%)and the isocortex (23.55% ± 14.66%). Finally, similar to LS-vmH neurons, LS-VTA neurons also receive the greatest proportion of input from the hippocampal formation (31.09% ± 7.21%) and the isocortex (27.59% ± 8.51%) (Figure 3E; two-way ANOVA with projection population and major brain region as factors, interaction: p = 0.0028, projection population: p = 0.998, major brain region: p = 1.01*10^-28^ with post-hoc t tests, see Supplementary Table 1). Therefore, LS neurons projecting to the PAG receive the most distinct pattern of input from major brain regions. Once we determined the largest sources of input to each projection population, we then chose to examine the inputs from each major input region separately using the high spatial resolution afforded to us by our Wholebrain mapping strategy.

**Figure 3.**
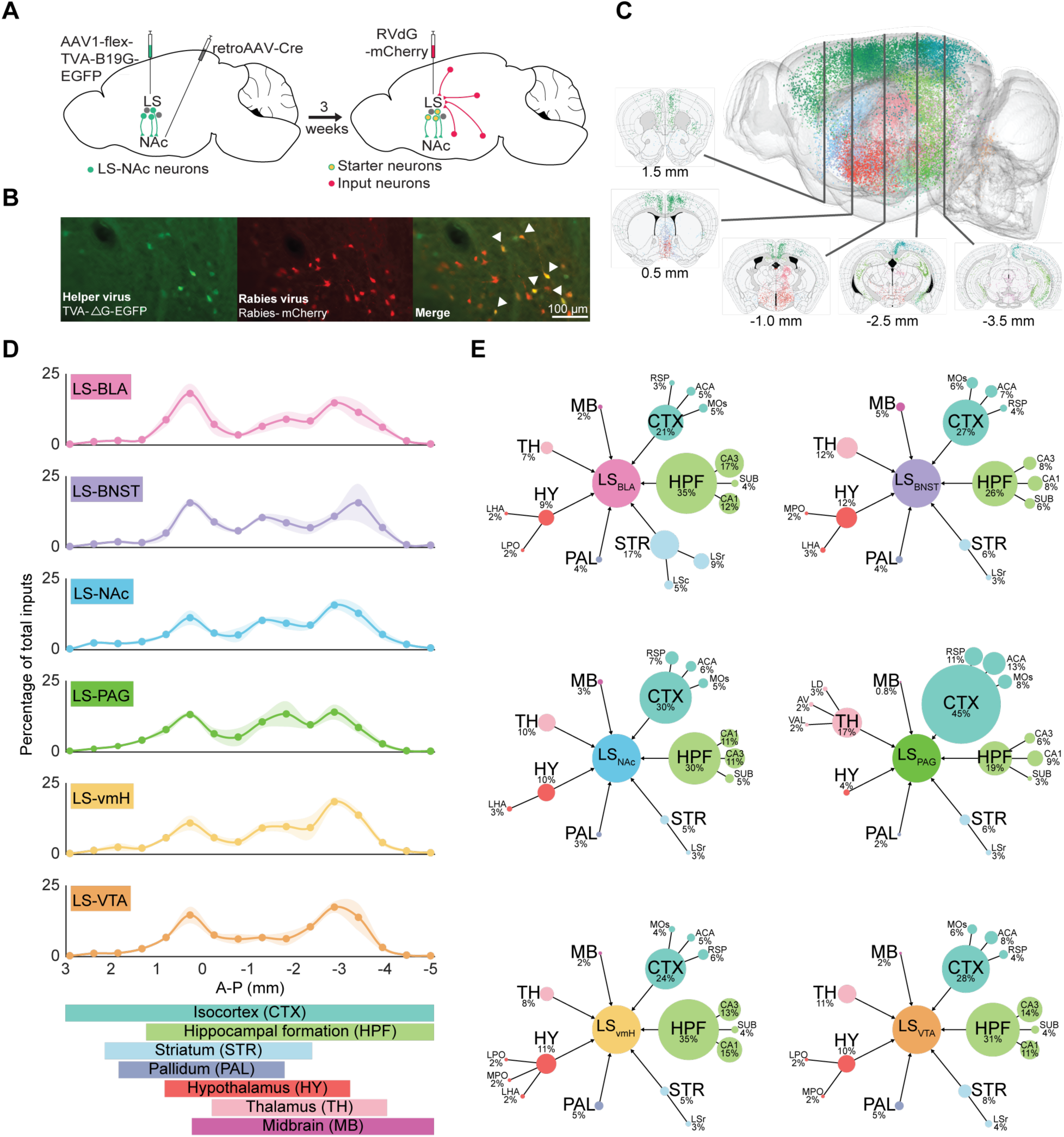
The LS-PAG projection population receives different proportions of inputs from major brain regions. A) Schematic of the viral tracing strategy used to label monosynaptic inputs to specific LS projection populations in mCherry, example shown is LS-NAc. The LS projection population is first retrogradely labeled with a helper virus tagged with GFP (left schematic). After three weeks, a modified rabies virus tagged with mCherry (RVdG-mCherry) is injected into the LS, labeling whole brain monosynaptic inputs to the GFP-tagged LS projection population with mCherry (right schematic). B) Example histology of starter cells in the LS at 20x magnification. Starter cells are neurons that have been labeled with both the helper virus and rabies virus. Cells labeled with the GFP-tagged helper virus are shown in green (left panel, green channel), cells labeled with the mCherry-tagged rabies virus are shown in red (middle panel, red channel), starter cells labeled with both GFP and mCherry are shown in yellow and indicated by filled arrowheads (right panel, overlay). Scale bar represents 100 μm. C) A three-dimensional representation of all monosynaptic input neurons to the LS-NAc projection population from a representative animal. Insets show input neurons mapped at individual A-P coordinates on coronal sections. Each dot represents one neuron. Dot colors correspond to different input regions as defined by the Allen Brain Atlas (isocortex: teal, hippocampal formation: green, striatum: light blue, pallidum: blue gray, hypothalamus: red, thalamus: pink, and midbrain: magenta). D) The percentages of brainwide inputs to each projection population along the entire A-P axis of the brain. Brain regions corresponding to a particular A-P range are listed below. Two-way ANOVA with projection population and bin as factors (interaction: p = 0.878, projection population: p = 1, bin: p = 3.43*10^-53^) with post-hoc t tests (see Supplementary Table 1). E) The percentage of inputs to LS projection populations from the major brain regions are similar except for the LS-PAG population, with the HPF being the greatest input for most of the LS projections. Within the LS-BLA projection, the HPF is a significantly larger input than the TH, HYP, MB and PAL. Within the LS-BNST projection, the CTX and HPF send more input than the MB and PAL. Within the LS-NAc, LS-vmH and LS-VTA projections, the HPF is a greater input than all major brain regions except CTX. CTX is a greater input to LS-vmH than MB and to LS-VTA than MB and PAL. In contrast, the LS-PAG population receives more input from CTX than any other major brain region, and the CTX sends a greater percentage of inputs to the LS-PAG population than the LS-BLA and LS-vmH projection populations. Two-way ANOVA with projection population and major regions as factors (interaction: p = 0.003, projection population: p = 0.998, major region: p = 1.01*10^-28^) with post-hoc t tests (see Supplementary Table 1). Average percentages were calculated by normalizing the raw count of neurons in each major region (hippocampal formation, isocortex, striatum, pallidum, hypothalamus, thalamus, midbrain) to the total number of input neurons per animal then calculating the mean percentage per projection population. Subregions were included in the spider map when they comprised at least 2% of the total inputs to the LS projection population. Circle areas correspond to the calculated percentage and are scaled consistently across spider maps. Subregion abbreviations: anterior cingulate area (ACA), anteroventral nucleus of thalamus (AV), lateral dorsal nucleus of thalamus (LD), lateral hypothalamic area (LHA), lateral preoptic area (LPO), caudal lateral septum (LSc), rostral lateral septum (LSr), secondary motor area (MOs), medial preoptic area (MPO), retrosplenial area (RSP), subiculum (SUB), ventral anterior-lateral complex of the thalamus (VAL). Shaded error regions indicate ± SEM. Representative histology shown in B is from the LS-vmH projection population. N = 3 animals per projection population.

### Varied isocortical inputs to LS projection populations

The isocortex as a whole sends proportionally large inputs to all of the LS projection populations. However, the isocortex, as defined by the Allen Brain Atlas, encompasses a vast number of disparate cortical regions, so we divided the isocortical inputs into anterior (A-P coordinates: 2.545 mm anterior to bregma to 1.055 mm posterior to bregma), posterior (A-P coordinates: 1.155 mm posterior to bregma to 4.955 mm posterior to bregma), and sensorimotor (A-P coordinates: 2.545 mm anterior to bregma to 4.955 mm posterior to bregma) isocortex. In the anterior isocortex (Figure 4A), all LS projection populations receive the greatest isocortical input from the anterior cingulate area (ACA), except for the LS-vmH, which receives a similar input from the infralimbic area. Additionally, compared to all of the other projection populations, the LS-PAG receives the greatest proportion of ACA input (% ACA input by projection population - LS-BLA: 5.19% ± 1.28%, LS-BNST: 6.92% ± 2.41%, LS-NAc: 5.91% ± 0.76%, LS-PAG: 12.97% ± 2.21%, LS-vmH: 4.68% ± 3.65%, LS-VTA: 7.56% ± 2.21%; two-way ANOVA with projection population and anterior isocortical region as factors, interaction: p = 2.25*10^-5^, projection population: p = 3.79*10^-4^, region: p = 3.25*10^-23^ with post-hoc t tests, see Supplementary Table 1). In the posterior isocortex (Figure 4B), the retrosplenial cortex (RSP) provides proportionally greater inputs than the parietal cortex to all LS projection populations and the LS-BLA receives less posterior isocortical input than the LS-PAG (% RSP input by projection population - LS-BLA: 2.82% ± 1.59%, LS-BNST: 4.20% ± 1.41%, LS-NAc: 7.49% ± 2.86%, LS-PAG: 10.59% ± 4.30%, LS-vmH: 6.25% ± 5.34%, LS-VTA: 4.90% ± 1.76%; two-way ANOVA with projection population and posterior isocortical region as factors, interaction: p = 0.321, projection population: p = 0.040, region: p = 4.67*10^-5^ with post-hoc t tests, LS-BLA to LS-PAG: p = 0.021, RSP to PTL: p = 4.67*10^-5^). In the sensorimotor isocortex (Figure 4C), the secondary motor area (MOs) provides a greater proportion of input to all LS projection populations than the somatosensory and primary motor areas (% MOs input by projection population - LS-BLA: 5.07% ± 3.31%, LS-BNST: 6.36% ± 3.86%, LS-NAc: 5.37% ± 1.74%, LS-PAG: 8.26% ± 3.43%, LS-vmH: 3.57% ± 2.89%, LS-VTA: 5.73% ± 2.10%; two-way ANOVA with projection population and sensorimotor isocortical region as factors, interaction: p = 0.938, projection population: p = 0.263, region: p = 2.36*10^-7^ with post-hoc t tests, MOs to MOp: p = 8.63*10^-7^, MOs to SS: p = 5.57*10^-6^). Consequently, there are large isocortical inputs into all LS projection populations, but the extent to which these inputs innervate different LS projection populations varies by isocortical region, especially the ACA.

**Figure 4.**
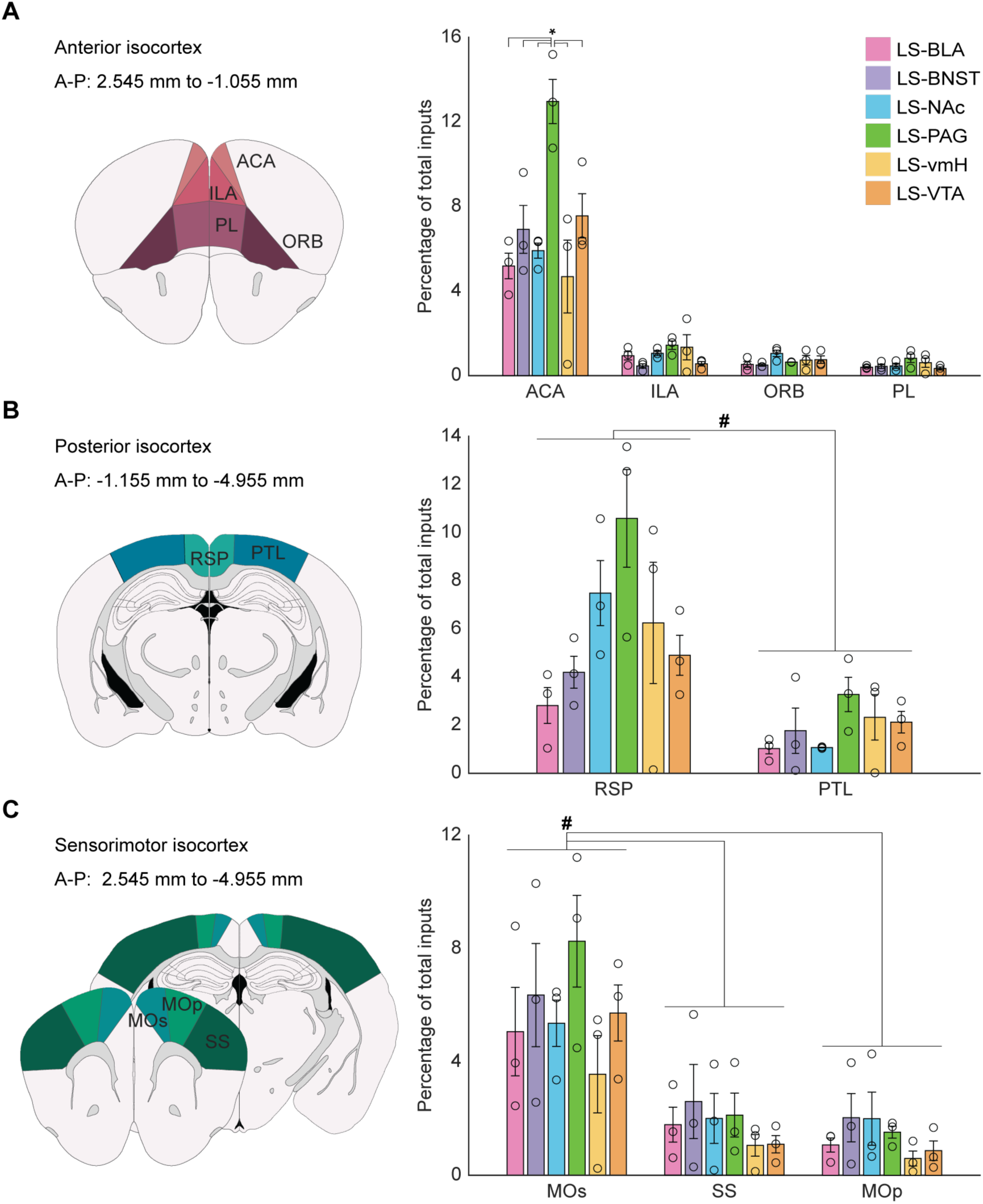
Isocortical inputs to individual LS projection populations vary by region. A) The anterior cingulate area (ACA) sends a significantly greater proportion of input to the LS-BLA, LS-BNST, LS-NAc, LS-PAG, and LS-VTA populations compared to the rest of the anterior isocortex. Of the ACA inputs, the LS-PAG population receives a larger proportion of ACA input than the other LS projection populations. Two-way ANOVA with projection population and region (ACA, infralimbic area (ILA), orbital area (ORB), prelimbic area (PL)) as factors (interaction: p = 2.25*10^-5^, projection population: p = 3.79*10^-4^, region: p = 3.25*10^-23^) with post-hoc t tests comparing projection population within region (ACA -LS-PAG to LS-BLA: p = 6.33*10^-8^, LS-PAG to LS-BNST: p = 3.03*10^-5^, LS-PAG to LS-NAc: p = 8.42*10^-7^, LS-PAG to LS-vmH: p = 1.07*10^-8^, LS-PAG to LS-VTA: p = 2.88*10^-4^) and region within projection population (see Supplementary Table 1). B) The retrosplenial area (RSP) has significantly more input to the LS projection populations than the posterior parietal association areas (PTL), and the LS-PAG received significantly more posterior isocortical input than the LS-BLA population. Two-way ANOVA with projection population and region (RSP and PTL) as factors (interaction: p = 0.321, projection population: p = 0.041, region: p = 4.67*10^-5^) with a post-hoc t test comparing regions (p = 4.67*10^-5^) and projections (LS-BLA to LS-PAG: p = 0.021). C) The secondary motor area (MOs) contributes significantly more input to the LS projection populations than the somatosensory areas (SS) or the primary motor area (MOp). There was no significant effect of projection population on sensorimotor inputs. Two-way ANOVA with projection population and region (MOs, SS, MOp) as factors (interaction: p = 0.938, projection population: p = 0.263, region: p = 2.36*10^-7^) with post-hoc t tests comparing regions (MOs to MOp: p = 8.63*10^-7^, MOs to SS: p = 5.57*10^-6^).Values from individual animals indicated by unfilled circles. Error bars indicate ± SEM. Colors correspond to the legend in A. *p < 0.05 for t tests following significant interaction; #p < 0.5 for t tests following significant main effect. N = 3 animals per projection population.

### A dorsoventral gradient of hippocampal inputs to LS projection populations

The LS has long been described as the major output region of the hippocampus. Historically, these inputs from the hippocampus have been described as topographically organized, such that the dorsal LS receives largely dorsal hippocampal input and the ventral LS receives largely ventral hippocampal input^4,13^. However, it is unknown if this topographical organization applies to individual regions of the hippocampus, such as CA1 and CA3. In corroboration with the aforementioned anatomical studies, we found that the hippocampal formation, along with the isocortex, provides one of the largest inputs (∼20-30% of the total inputs) to each LS projection population, except for the LS-PAG which receives a larger isocortical input (Figure 3E). We then quantified the proportion of inputs from separate hippocampal areas and found that these populations receive proportionally similar inputs across hippocampal subregions (Figure 5A and B; one-way ANOVA, CA3: p = 0.249, CA1: p = 0.209, SUB: p = 0.698, CA2: p = 0.560, DG: p = 0.213, ENT: p = 0.210).

**Figure 5.**
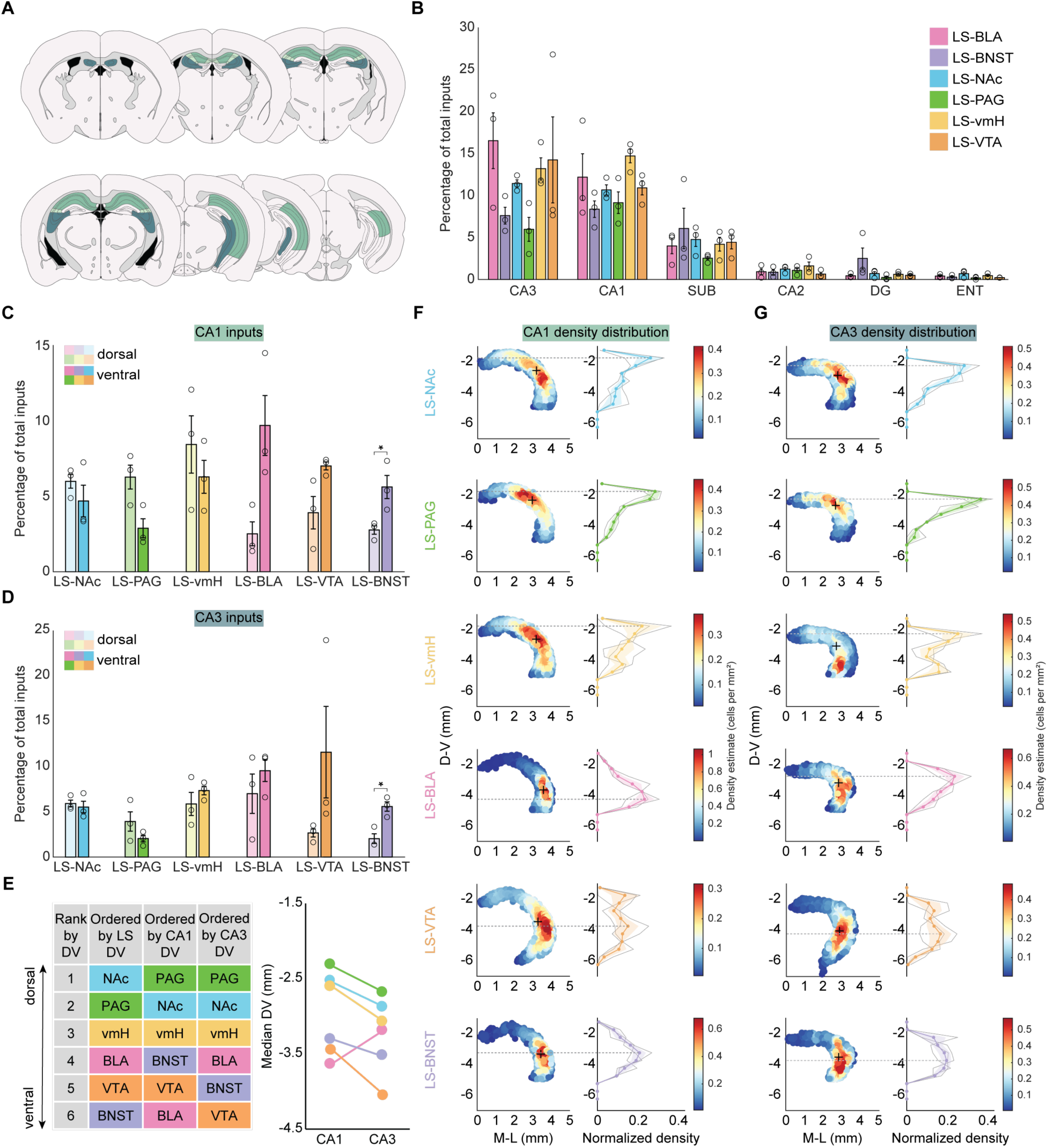
Hippocampal inputs to LS projection populations are topographically organized in a dorsoventral gradient. A) Schematic showing coronal sections spanning the A-P axis of the hippocampus, which contains the hippocampal areas: CA1 (green), CA2 (light green), and CA3 (blue). B) Each LS projection population receives a similar percentage of inputs from various subregions of the hippocampal formation (CA3, CA1, subiculum (SUB), CA2, dentate gyrus (DG), entorhinal area (ENT)). One-way ANOVA (CA3: p = 0.249, CA1: p = 0.209, SUB: p = 0.698, CA2: p = 0.560, DG: p = 0.213, ENT: p = 0.210). C) When CA1 inputs are classified as dorsal (light) or ventral (dark) using a D-V coordinate cutoff of −3 mm, the LS-BNST projection population receives significantly more ventral CA1 than dorsal CA1 input. Unpaired t test (LS-NAc: p = 0.404, LS-PAG: p = 0.052, LS-vmH: p = 0.469, LS-BLA: p = 0.052, LS-VTA: p = 0.083, LS-BNST: p = 0.047). D) When CA3 inputs are classified as dorsal (light) or ventral (dark) using a D-V coordinate cutoff of −3 mm on the D-V axis, the LS-BNST projection population receives significantly more ventral CA3 than dorsal CA3 input. Unpaired t test (LS-NAc: p = 0.696, LS-PAG: p = 0.244, LS-vmH: p = 0.424, LS-BLA: p = 0.451, LS-VTA: p = 0.225, LS-BNST: p = 0.018). E) Ranking of LS projection populations, CA1 inputs, and CA3 inputs by median D-V coordinate demonstrate a dorsoventral gradient of hippocampal inputs to LS projection populations, with more dorsally located LS populations receiving more dorsal CA1 and CA3 inputs and more ventrally location LS populations receiving more ventral CA1 and CA3 inputs (left). With the exception of the LS-BLA population, the ordering of the median D-V coordinates of the CA1 and CA3 inputs are preserved across the LS projection populations. F) Density plots (left column) and distribution of density (right column) of CA1 inputs along the D-V axis of the hippocampus (A-P coordinate: −1.355 to −3.78 mm) shows topographically organized inputs to LS projection populations. The + indicates the median M-L and D-V coordinate for the CA1 inputs to each LS projection population (+ M-L/D-V coordinates - LS-NAc: 3.20/−2.54, LS-PAG: 2.98/−2.33, LS-vmH: 3.20/−2.61, LS-BLA: 3.59/−3.64, LS-VTA: 3.30/−3.44, LS-BNST: 3.45/−3.31). The dashed gray line spanning the left and right panels corresponds to the peak of the normalized density distribution plot (D-V of gray line - LS-NAc: −1.75 mm, LS-PAG: −1.75 mm, LS-vmH: −1.75 mm, LS-BLA: −4.25 mm, LS-VTA: −3.75 mm, LS-BNST: −3.25 mm) G) Density plots (left column) and distribution of density (right column) of CA3 inputs along the D-V axis of the hippocampus (A-P coordinate: −0.955 mm to −3.455 mm) shows relative topographical organization of inputs to LS projection populations. The + indicates the median M-L and D-V coordinate for the CA3 inputs to each LS projection population (+ M- L/D-V coordinates - LS-NAc: 2.77/−2.88, LS-PAG: 2.65/−2.69, LS-vmH: 2.70/−3.07, LS- BLA: 2.83/−3.20, LS-VTA: 2.84/−4.04, LS-BNST: 2.83/−3.53). The dashed gray line spanning the left and right panels corresponds to the peak of the normalized density distribution plot (D-V of gray line - LS-NAc: −2.25 mm, LS-PAG: −2.25 mm, LS-vmH: −2.25 mm, LS-BLA: −2.75 mm, LS-VTA: −4.25 mm, LS-BNST: −3.75 mm). The LS projection populations are ordered from dorsal to ventral by the median D-V coordinate of each population quantified using the retrograde tracing strategy in Figure 1G. Coordinates are relative to bregma. Colors correspond to the legends in B and C. Values from individual animals indicated by unfilled circles or gray lines. Error bars and shaded error regions indicate ± SEM. *p < 0.05. N = 3 animals per projection population.

Due to our previous identification of a dorsoventral organization of projection populations within the LS (Figure 1G; Figure 2), we were next able to determine if the topographical organization of hippocampal inputs is maintained across these six anatomically organized projection populations. In order to do this, we classified CA1 and CA3 inputs as either dorsal or ventral based on a D-V coordinate cutoff of −3 mm. We then quantified the proportion of total inputs that arose from dorsal (CA1 - LS-NAc: 5.99% ± 0.975%, LS-PAG: 6.27% ± 1.68%, LS-vmH: 8.44% ± 4.03%, LS-BLA: 2.51% ± 1.68%, LS-VTA: 3.91% ± 2.28%, LS-BNST: 2.76% ± 0.597%; CA3 - LS-NAc: 5.91% ± 0.802%, LS-PAG: 3.93% ± 2.27%, LS-vmH: 5.87% ± 2.70%, LS-BLA: 7.00% ± 4.62%, LS-VTA: 2.67% ± 0.940%, LS-BNST: 2.04% ± 1.14%) and ventral (CA1 - LS-NAc: 4.69% ± 2.22%, LS-PAG: 2.88% ± 1.34%, LS-vmH: 6.29% ± 2.32%, LS-BLA: 9.70% ± 4.22%, LS-VTA: 7.01% ± 0.564%, LS-BNST: 5.62% ± 1.63%; CA3 - LS-NAc: 5.53% ± 1.35%, LS-PAG: 2.04% ± 0.777%, LS-vmH: 7.35% ± 0.997%, LS-BLA: 9.54% ± 2.54%, LS-VTA: 11.59% ± 10.74%, LS-BNST: 5.58% ± 1.10%) CA1 and CA3 regions and sorted them based on the D-V organization of LS projection populations (Figure 5C and D). We found that the most ventrally residing LS projection population, the LS-BNST, received significantly more ventral than dorsal CA1 and CA3 inputs, while the rest of the projection populations received proportionally similar dorsal and ventral inputs from both hippocampal areas. (Unpaired t test; CA1 - LS-NAc: p = 0.404, LS-PAG: p = 0.052, LS-vmH: p = 0.469, LS-BLA: p = 0.052, LS-VTA: p = 0.084, LS-BNST: p = 0.047; CA3 - LS-NAc: p = 0.696, LS-PAG: p = 0.244, LS-vmH: p = 0.424, LS-BLA: p = 0.451, LS-VTA: p = 0.225, LS-BNST: p = 0.018).

With the slight topographical organization detected by examining proportional inputs, we utilized the high spatial resolution of the Wholebrain input mapping to plot the density distribution of each CA1 and CA3 neuron across all animals along the A-P axis of the hippocampal formation (Figure 5F and G). Using CA1 and CA3 density distributions, a topographical organization of hippocampal inputs emerges, such that the median D-V coordinate of the distribution of inputs from CA1 and CA3 tends to decrease along the D-V gradient of LS projection populations (CA1 median D-V coordinate (mm) - LS-NAc: −2.54, LS-PAG: −2.33, LS-vmH: −2.61, LS-BLA: −3.64, LS-VTA: −3.44, LS-BNST: −3.31; CA3 median D-V coordinate (mm) - LS-NAc: −2.88, LS-PAG: −2.69, LS-vmH: −3.07, LS-BLA: −3.20, LS-VTA: −4.04, LS-BNST: −3.53). As a result, we show a correlation between the D-V location of the projection population within the LS and the D-V location within the hippocampus of the CA1 and CA3 input it receives (Figure 5E). For example, LS-NAc and LS-PAG are the most dorsal LS populations, and they receive the highest density of CA1 and CA3 inputs from a more dorsal portion of the hippocampus, while the LS-vmH is an intermediate LS population that receives CA1 and CA3 input from intermediate portions of the hippocampus (Figure 5E-G). The LS-BNST is a ventral LS population that receives a high density of inputs from ventral hippocampal regions (Figure 5E-G). The LS-VTA receives slightly more ventral hippocampal inputs compared to where it is located in the LS. The LS-BLA is an intermediate LS population that receives intermediate CA3 input but the most ventral CA1 input. Consequently, there are slight variations to the topographical organization of hippocampal inputs to LS projection populations depending on the hippocampal area. The hippocampal area CA2 (Figure 5A) has an extensive role in social behaviors^25,51,52^. However, the analysis of the topographical organization of CA2 inputs into the LS is limited because the Allen Brain Atlas does not include a ventral CA2 region. We find that all six LS projection populations receive a similar proportion of CA2 inputs (Supplementary Figure 5A; one-way ANOVA, CA2: p = 0.560) and that these CA2 inputs are similarly distributed (Supplementary Figure 5B), again limited by the current divisions of the Allen Brain Atlas. Overall, we are able to recapitulate and expand upon the findings of previous anatomical studies regarding the topographical organization of hippocampal input to discrete LS projection populations depending on the hippocampal area from which the input originates.

### Individual subcortical brain regions differ in specificity of inputs to each projection population

Along with cortical and hippocampal inputs, the LS receives inputs from subcortical structures. Unlike the hippocampal inputs to the LS, little is known about how the subcortical inputs to the LS are organized. These inputs are proportionally smaller than the cortical and hippocampal inputs but have been proposed to provide information about internal state, which may be integrated in the LS alongside contextual information from the hippocampus and cortex^8,9^. Two subcortical regions that each contribute approximately 10% of the total input to each projection population (Figure 3E) are the thalamus (% thalamic input by projection population - LS-BLA: 7.03% ± 2.14%, LS-BNST: 11.86% ± 4.44%, LS-NAc: 10.04% ± 3.27%, LS-PAG: 16.99% ± 4.44%, LS-vmH: 8.20% ± 4.91%, LS-VTA: 10.71% ± 1.66%) and hypothalamus (% hypothalamic input by projection population - LS-BLA: 9.13% ± 1.49%, LS-BNST: 12.11% ± 5.54%, LS-NAc: 9.81% ± 2.14%, LS-PAG: 3.55% ± 1.35%, LS-vmH: 11.66% ± 10.37%, LS-VTA: 9.90% ± 8.22%).

We visualized the density of cell counts across all animals at three different ranges of A-P coordinates of the thalamus (Figure 6A). We identified clusters of neurons across the three A-P coordinates that corresponded to various thalamic nuclei, so we then ranked the top ten thalamic inputs by projection population (Figure 6B). Some LS projection populations had similar rankings of thalamic input, such as the LS-BNST, LS-PAG and LS-vmH, which all received the most input from the lateral dorsal nucleus of the thalamus (LD) (Figure 6B). Whereas, other LS projection populations received the most input from different thalamic nuclei, such as the LS-BLA which received the most input from the ventral anterior-lateral complex of the thalamus (VAL) and the LS-VTA which received the most input from the anteromedial nucleus (AM) (Figure 6B). The LS-NAc received similar proportions of input from two thalamic nuclei, the AM and LD nuclei (Figure 6B). In addition, we found that thalamic inputs vary in terms of which nuclei project to each projection population, with some nuclei projecting robustly to all LS populations (Figure 6C, D) and others exclusively projecting to some LS populations but not others (Figure 6E-H).

**Figure 6.**
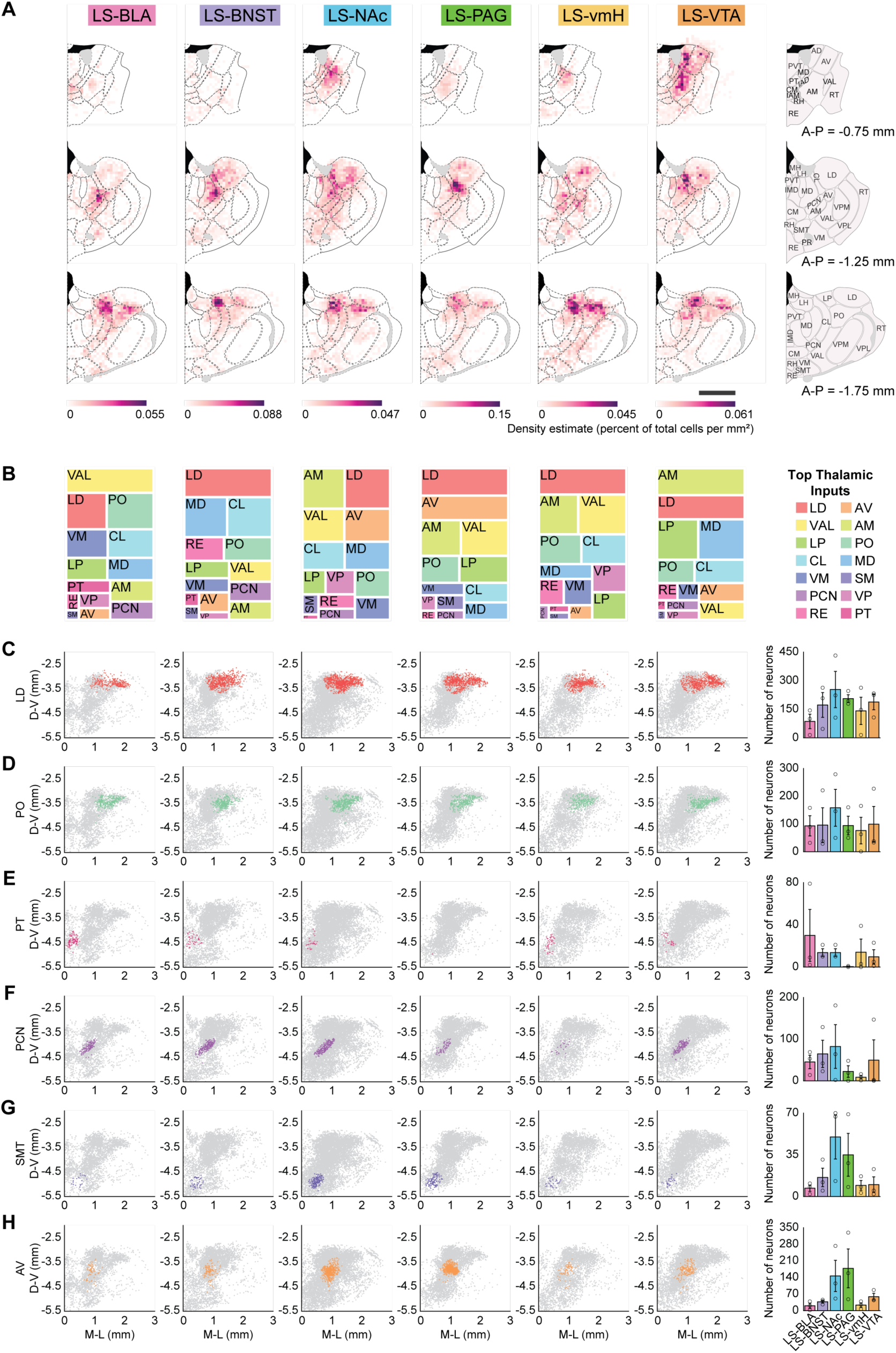
Thalamic nuclei specifically innervate different LS projection populations. A) Density estimates of thalamic inputs to the LS projection populations collapsed across three different ranges of A-P coordinates: −0.48 mm to −0.9 mm (top row), −0.9 mm to −1.455 mm (middle row), and −1.455 mm to −2.155 mm (bottom row) with the corresponding map of thalamic nuclei from coronal sections of the thalamus at each A-P coordinate (right panel). B) Rank ordering of the top thalamic inputs across all LS projection populations. The area of each region on the treemap corresponds to the proportional contribution (average normalized input) of a region relative to all other top thalamic nuclei. C) Inputs from the lateral dorsal nucleus of the thalamus (LD) across all animals from each projection population. Average LD input neuron counts across projection populations (right panel). D) Inputs from the posterior complex of the thalamus (PO) across all animals from each projection population. Average PO input neuron counts across projection populations (right panel). E) Inputs from the parataenial nucleus (PT) across all animals from each projection population. Average PT input neuron counts across projection populations (right panel). F) Inputs from the paracentral nucleus (PCN) across all animals from each projection population. Average PCN input neuron counts across projection populations (right panel). G) Inputs from the submedial nucleus of the thalamus (SMT) across all animals from each projection population. SMT input neuron counts across projection populations (right panel). H) Inputs from the anteroventral nucleus of the thalamus (AV) across all animals from each projection population. Average AV counts across projection populations (right panel). Thalamic nuclei abbreviations: Anterodorsal nucleus (AD), Anteromedial nucleus (AM), Anteroventral nucleus of thalamus (AV), Central lateral nucleus of thalamus (CL), Central medial nucleus of the thalamus (CM), Interanterodorsal nucleus of the thalamus (IAD), Interanteromedial nucleus of the thalamus (IAM), Intermediodorsal nucleus of the thalamus (IMD), Lateral dorsal nucleus of thalamus (LD), Lateral habenula (LH), Lateral posterior nucleus of the thalamus (LP), Mediodorsal nucleus of thalamus (MD), Medial habenula (MH), Paracentral nucleus (PCN), Posterior complex of the thalamus (PO), Perireunensis nucleus (PR), Parataenial nucleus (PT), Paraventricular nucleus of the thalamus (PVT), Nucleus of reuniens (RE), Rhomboid nucleus (RH), Reticular nucleus of the thalamus (RT), Submedial nucleus of the thalamus (SMT), Ventral anterior-lateral complex of the thalamus (VAL), Ventral medial nucleus of the thalamus (VM), Ventral posterolateral nucleus of the thalamus (VPL), Ventral posteromedial nucleus of the thalamus (VPM). Each colored dot represents an input neuron from a specific thalamic subregion and each gray dot represents a thalamic input neuron. Values from individual animals indicated by unfilled circles. Error bars indicate ± SEM. N = 3 animals per projection population.

It was theorized that the hypothalamic inputs may also be topographically organized, similarly to the LS outputs to the hypothalamus which are organized such that dorsal LS projects to the lateral hypothalamus and progressively more ventral regions of the LS project to progressively more medial regions of the hypothalamus^4^. However, we did not find any clear organization of hypothalamic inputs to the LS, even along its M-L axis. In fact, the hypothalamic nuclei send largely uniform inputs to the LS projection populations, with the largest input coming from the lateral hypothalamus (LHA) across all projection populations, except for the LS-VTA (Figure 7A, number of LHA neurons by projection population - LS-BLA: 248.33 ± 19.14, LS-BNST: 338.67 ± 263.51, LS-NAc: 780.33 ± 517.01, LS-PAG: 90.33 ± 31.56, LS-vmH: 403.33 ± 386.58, LS-VTA: 254.33 ± 176.89). We found that with the exception of the LS-PAG population, hypothalamic subregions similarly innervate the other five LS projection populations studied (Figure 7B-F). It has also been suggested that reciprocal connections between the hypothalamus and LS may serve as a feedback loop to terminate ongoing social behaviors. We provide evidence of reciprocal connectivity between the LS and the vmH (Figure 7F), a putative mechanism by which the vmH itself might regulate ongoing aggressive behaviors. With the maps that we have provided of the subcortical inputs to the LS, additional work can be done to clarify how these inputs function, especially in social behaviors.

**Figure 7.**
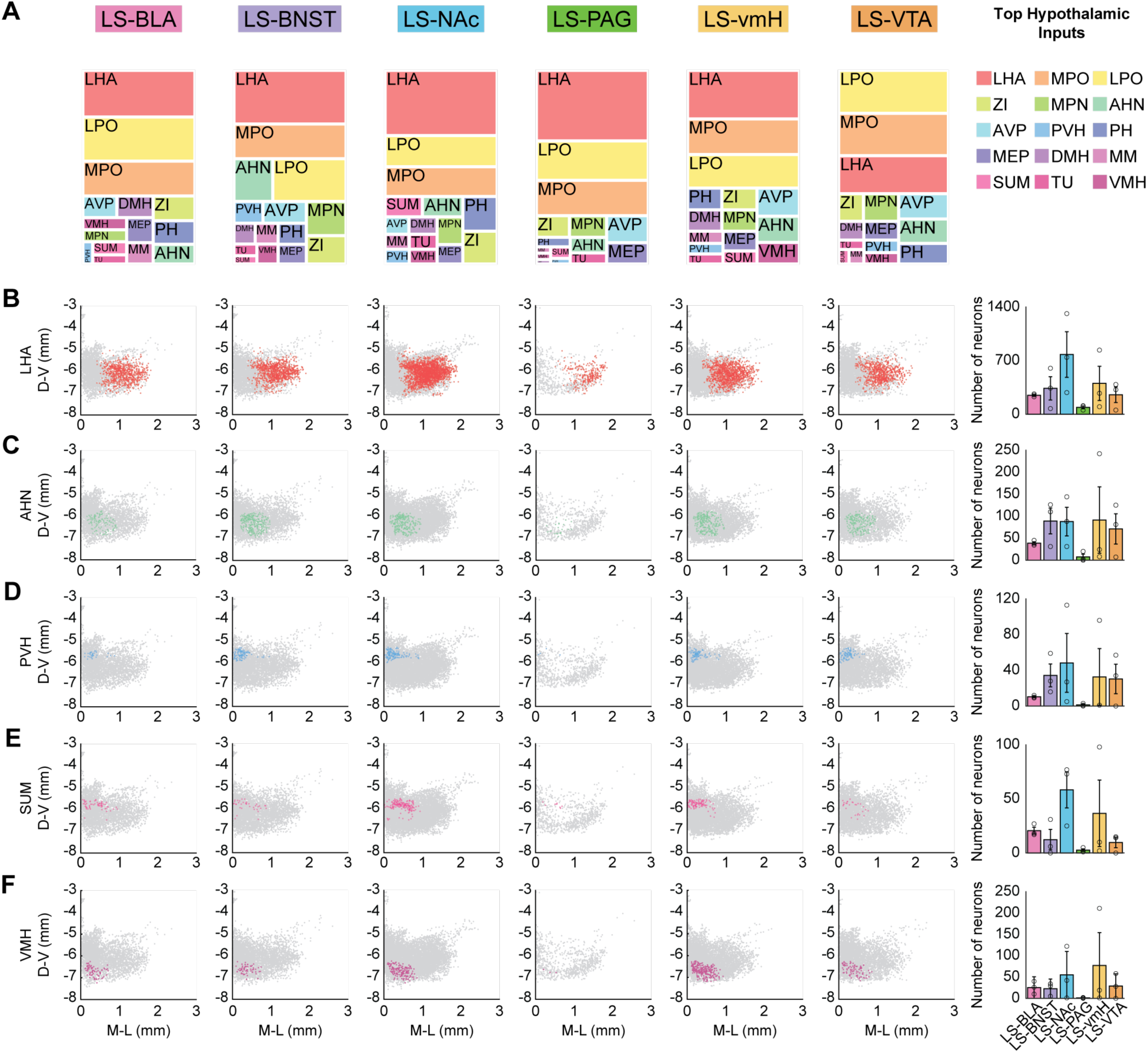
Hypothalamic inputs are similar across LS projection populations. A) Rank ordering of the top hypothalamic inputs across all LS projection populations. The area of each region on the treemap corresponds to the proportional contribution (average normalized input) of a region relative to all the other top thalamic nuclei. B) Inputs from the lateral hypothalamic area (LHA) across all animals from each projection population. Average LHA input neuron counts across projection populations (right panel). C) Inputs from the lateral hypothalamic area (AHN) across all animals from each projection population. Average AHN input neuron counts across projection populations (right panel). D) Inputs from the lateral hypothalamic area (PVH) across all animals from each projection population. Average PVH input neuron counts across projection populations (right panel). E) Inputs from the lateral hypothalamic area (SUM) across all animals from each projection population. Average SUM input neuron counts across projection populations (right panel). F) Inputs from the lateral hypothalamic area (VMH) across all animals from each projection population. Average VMH input neuron counts across projection populations (right panel). Hypothalamic subregion abbreviations: Anterior hypothalamic nucleus (AHN), Anteroventral preoptic nucleus (AVP), Dorsomedial nucleus of the hypothalamus (DMH), Lateral hypothalamic area (LHA), Lateral preoptic area (LPO), Median preoptic nucleus (MEP), Medial mammillary nucleus (MM), Medial preoptic nucleus (MPN), Medial preoptic area (MPO), Posterior hypothalamic nucleus (PH), Paraventricular hypothalamic nucleus (PVH), Supramammillary nucleus (SUM), Tuberal nucleus (TU), Ventromedial hypothalamic nucleus (VMH), zona incerta (ZI). Each colored dot represents an input neuron from a specific hypothalamic subregion and each gray dot represents a hypothalamic input neuron. Values from individual animals indicated by unfilled circles. Error bars indicate ± SEM. N = 3 animals per projection population.

## Discussion

We combined brainwide mapping of fluorescently labeled neurons with retrograde and monosynaptic viral tracing strategies to provide comprehensive high-resolution maps of six projection populations within the LS, as well as the inputs to each of these projection populations. We find that the LS projects to brain regions involved in regulating a variety of social behaviors and that projection populations are organized into discrete anatomical regions within the LS, especially along its dorsoventral axis. Additionally, by mapping brainwide inputs to each of these projection populations, we identify unique patterns of innervation in a region-dependent manner.

### Anatomical mapping of LS connectivity

The LS occupies an anatomically significant location in the brain. It receives large hippocampal and cortical inputs and directs these inputs to many downstream subcortical structures. While the existing literature has provided valuable insight into the anatomy of the LS itself and its inputs from the hippocampus, less is known about its other sources of inputs^4,12^. Additionally, these studies had predominantly focused on LS outputs to the hypothalamus. Since then, novel cell-type specific sequencing and spatial profiling has been performed within the LS to corroborate and expand upon findings from these early anatomical studies^19–21^. However, large-scale efforts to systematically map the brainwide anatomical connectivity of the LS using modern viral tracing strategies had not yet been performed. In this study, we use high-resolution brainwide mapping techniques to characterize inputs across a wide range of brain regions onto specific LS projection populations. Thus, we are able to describe how inputs to various LS populations are organized.

### Organization of projection populations in the LS

In earlier anatomical studies of the LS, it was suggested that the topographical organization of LS inputs to the hypothalamus corresponded with different behavioral outputs mediated by different hypothalamic regions^4^. In particular, it was proposed that the caudal LS (LSc) which receives inputs from neuromodulatory centers in the brainstem, as well as from the lateral hypothalamus, and projects to the medial septum and supramammillary nucleus was involved in theta rhythm generation related to locomotion^4,5^. In contrast, the rostral LS (LSr) which is reciprocally connected with other hypothalamic regions mediated reproductive and social behaviors^4^. It was also proposed that the ventral LS (LSv) which receives a large input from the ventral hippocampus and projects to more medial hypothalamic nuclei regulated ingestive and homeostatic behaviors^4^. However, the organization of LS neurons that project to non-hypothalamic brain regions is relatively poorly understood. We directly addressed this in our study by examining LS inputs to six different projection populations. We find that LS projections to brain regions outside of the hypothalamus occupy discrete regions within the LS and that they are organized along the anatomical axes of the LS (Figure 1E-H; Figure 2), especially along the dorsoventral axis (Figure 1G). This finding further supports the idea that distinct regions of the LS project to different downstream targets and mediate different behaviors^9^. Although there is evidence of organization of information along the dorsoventral axis of the LS^24,25^, we are among the first to provide evidence of anatomical organization of non-hypothalamic LS projections, which may serve as a substrate by which this functional information is organized.

### Organization of cortical and hippocampal inputs to the LS

Different theories of LS function posit that the organization of inputs that it receives will determine how it transmits information to downstream structures^9^. Here, we provide detailed maps of cortical and hippocampal inputs to discrete LS projection populations. In particular, we find that the LS-PAG projection population receives more isocortical input, especially from the anterior cingulate area (Figure 4A), than any other projection population. This may suggest a unique role for LS-PAG neurons, similar to what has been found in cortical neurons projecting to the PAG^53^. Additionally, different hippocampal areas send topographically organized inputs to projection populations that are located in different regions along the dorsoventral axis of the LS. For example, the LS-BNST is the most ventral LS projection population and it receives a great proportion of ventral inputs from hippocampal areas CA1 and CA3 (Figure 5C and D). Whereas the two more dorsal LS projection populations, LS-NAc and LS-PAG receive a higher density of dorsal hippocampal input from CA1 and CA3 (Figure 5E-G). There is evidence of functional differences between dorsal and ventral regions of the hippocampus^54–56^. Thus, our findings demonstrate how distinct streams of cortical and hippocampal information may be transmitted through discrete LS neural populations to specific downstream targets to mediate different behaviors.

### Organization of subcortical inputs to the LS

Although there has been much emphasis on the cortical and hippocampal inputs to the LS, it also receives inputs from subcortical regions such as the thalamus and hypothalamus. As a result, we examined the patterns of input from these two subcortical regions. In the thalamus, we found specificity in how different thalamic nuclei project to neural populations within the LS (Figure 6). However, in the hypothalamus, there was not a clear anatomical organization of inputs into different LS projection populations, as was initially hypothesized^4^. We did identify reciprocal connections between a hypothalamic nucleus, the vmH, and the LS neurons that project to it (Figure 7F). This has interesting implications for how individual hypothalamic nuclei may provide feedback to the LS projections it receives to regulate certain behaviors, such as aggression.

### Limitations of the study

The Wholebrain mapping software we used to create both regional and brainwide maps of LS projections populations and their monosynaptic inputs employs coordinate systems and anatomical boundaries as determined by the Allen Brain Atlas. While this allows us to systematically map tissue sections across the entire brain and for consistent comparisons across animals, we are constrained by the anatomical boundaries imposed by the Allen Brain Atlas. In addition, as with most anatomical studies, we have to contend with the inherent anatomical variability in brains of individual animals. Furthermore, manually curating brainwide inputs to six individual LS projection populations across individual animals is a time and labor intensive process which limits the number of animals that can be included in our analysis.

### Broader implications

The LS is an important brain region involved in many motivated behaviors, including a diverse range of social behaviors. It is often treated as a homogeneous structure, despite evidence that it contains spatially organized and heterogeneous subregions. Consequently, results regarding its function are often conflicting^6^. Recently, efforts have been made to characterize the contribution of specific LS cell types to distinct behaviors. However, even these results do not provide a clear explanation of LS function^8^. Here, we examined LS projections to six different downstream regions as a putative substrate by which the LS may mediate distinct social behaviors. We found that these projection populations occupy discrete regions within the LS. In fact, the dorsoventral organization of discrete LS projection populations may correspond to previously identified LS cell types and subpopulations. It would be interesting to compare our maps of LS projection populations to those of cell type markers (Sst, Nts, and Esr1) and more detailed spatial transcriptomic maps of LS gene expression. This would allow us to further classify the identity of these projection population neurons. Additionally, the existence of different cell types within each projection population may help to explain the often conflicting results that arise from cell type or region specific manipulations. Further studies isolating specific cell types within a projection population may help to resolve how LS subpopulations contribute to different social behaviors. Finally, by providing detailed brainwide connectivity maps of individual LS projection populations, we hope to clarify how the anatomical organization of the LS may inform its function.

## Methods

### Experimental subjects

For all experiments, mice aged 8-12 weeks from Jackson laboratories were used. All experimental procedures were approved by the Emory Institutional Animal Care and Use Committee.

### Viruses

All viruses used in this study, except for the modified rabies virus, were purchased from Addgene: AAV5-pCAG-FLEX-EGFP-WPRE (51502-AAV5), AAVrg-EF1a-mCherry-IRES-Cre (55632-AAVrg), AAV1-synP-FLEX-splitTVA-EGFP-B19G (52473-AAV1), and pENN-AAVrg -hSyn-Cre-WPRE-hGH(105553-AAVrg). The modified rabies virus, Rabies-N2C-deletedG -mCherry-EnvA, was purchased from Thomas Jefferson University (Dr. Christoph Wirblich).

### Stereotaxic surgeries

For retrograde tracing surgeries, male mice aged ∼8-10 weeks (n = 18 mice, 3 per projection population) were anesthetized with 1-2% isoflurane and placed in a Kopf stereotactic system. A Nanoject microsyringe was used to inject 0.75 μL of AAV5-pCAG-FLEX-EGFP-WPRE (Addgene, injected titer of 1.3*10^13^ parts/mL) unilaterally into the LS (0 mm anterior, 0.4 mm lateral and 2.8 mm in depth) and 0.75 μL of AAVrg-EF1a-mCherry-IRES-Cre (Addgene, injected titer of 2.0*10^13^ parts/mL) into one of six ipsilateral target regions: the BLA (1.6 mm posterior, 3.4 mm lateral and 4.8 mm in depth); the BNST (0.15 mm anterior, 1.0 mm lateral and 4.3 mm in depth); the NAc (1.3 mm anterior, 1.3 mm lateral and 4.7 mm in depth); the PAG (4.6 mm posterior, 0.5 mm lateral and 2.2 mm in depth); the vmH (1.7 mm posterior, 0.68 mm lateral and 5.25 mm in depth); and the VTA (3.1 mm posterior, 0.35 mm lateral and 4.7 mm in depth). Mice were group housed and allowed to recover for three weeks following viral injection surgery to allow for sufficient viral expression, after which time they were sacrificed and their brain tissue collected for histological analysis.

For monosynaptic rabies tracing surgeries, male mice aged ∼8-10 weeks (n = 18 mice, 3 per projection population) were anesthetized with 1-2% isoflurane and placed in a Kopf stereotactic system. A Nanoject microsyringe was used to inject 0.75 μL of the Cre-dependent helper virus, AAV1-synP-FLEX-splitTVA-EGFP-B19G (Addgene, injected titer of 2.4*10^13^ parts/mL), unilaterally into the LS (0 mm anterior, 0.4 mm lateral and 2.8 mm in depth) and 0.75 μL of pENN-AAVrg-hSyn-Cre-WPRE-hGH (Addgene, injected titer of 1.9*10^13^ parts/mL) into one of six ipsilateral target regions: the BLA, BNST, NAc, PAG, vmH and VTA (same coordinates as retrograde tracing surgeries). Mice were group housed and allowed to recover for three weeks following viral injection surgery. At three weeks, mice were again anesthetized with 1-2% isoflurane and placed in a Kopf stereotactic system. A Nanoject microsyringe was used to inject 0.5 μL of the modified rabies virus, Rabies-N2C-deletedG-mCherry-EnvA (Thomas Jefferson University, injected titer of 8*10^8^ parts/mL), into the LS at the same location as the previous viral injection surgery. Mice were group housed and allowed to recover for seven days, after which they were sacrificed and their brain tissue collected for histological analysis.

### Histology and microscopy

Three weeks following retrograde injection surgeries and one week following rabies virus injection, mice were sacrificed and perfused with 0.5% PBS and 4% paraformaldehyde (PFA). The brains were dissected and fixed in 4% PFA overnight at 4°C. Brains were then transferred to 30% sucrose solution and allowed to fix for a minimum of 24 hours at 4°C. For all histological analysis, brains were sliced at 50 μm using an Epredia HM430 microtome and slices were mounted in DAPI-containing mounting solution (Product ID: 0100-20, Southern Biotech). For retrograde tracing experiments, every 50 μm slice containing the LS (from an approximate A-P of 1.3 mm anterior to 0.3 mm posterior) was mounted, as well as the slices containing the target brain region of interest. For monosynaptic rabies tracing experiments, every other 50 μm slice across the entire brain (from an approximate A-P of 2.545 mm to −4.955 mm) was mounted. Slices were then imaged using fluorescence microscopy (Keyence BZ-X800), and images were registered using Wholebrain software.

### Data analysis and statistics

#### Image processing and segmentation

After imaging the slides with a 4x objective lens, the individual sections were cropped and converted into 8-bit, gray-scaled TIFFs to be compatible with the Wholebrain R package used. Using Fiji from ImageJ, the color channels were split, and either the green channel (retrograde tracing) or red channel (modified rabies tracing) was saved. Rectangular ROIs were drawn around each individual slice on each slide, leaving a margin around the edge of the tissue. A custom macro was then used to crop each section based on its surrounding ROI, downsample the cropped TIFF into an 8-bit TIFF, then grayscale the resulting TIFF.

#### Wholebrain registration and cell-counting

An open-source Wholebrain R package was used to register each coronal brain section to a manually assigned anterior-posterior (A-P) coordinate based on the Allen Mouse Brain Reference Atlas. Each section was matched to a particular A-P coordinate using anatomical landmarks, such as the corpus callosum and hippocampal formation. Using Wholebrain, the individual section was registered to the designated A-P coordinate and mapped to the Allen Brain atlas as described in Fürth et al., 2018^50^. As part of its automated registration process, the Wholebrain software identifies the contour or edge of the tissue using a binary mask. Then, using the centroid of the identified contour and its resulting principal components, the software will generate correspondence points along the edge of the tissue. These correspondence points act as a reference from the tissue to the atlas section. Users can then manually add or remove correspondence points as needed to match the anatomy of the section of tissue to the atlas, and the thin-plate splines algorithm will run after each change. The registered image was then segmented using a soma size and brightness filter designed to identify individual neurons labeled by either a green (retrograde tracing) or red (monosynaptic rabies tracing) fluorophore. After individual neurons were detected by the software, neuron identity was manually confirmed for every section. In the case that the software failed to detect individual neurons, neurons were manually registered.

#### Quantification of LS neuron distributions

Analysis of the LS projection populations was restricted to cells labeled as belonging to the lateral septal complex (LSX) in the hemisphere ipsilateral to the viral injection, ranging from 1.3 mm to −0.3 mm on the A-P axis.

### Total number of LSX neurons

To quantify the number of neurons comprising each projection population (Figure 1D), we included all neurons that were identified as belonging to the LSX. These counts were averaged across each projection and plotted (± SEM) with individual animals. The one-way ANOVA was followed by post-hoc Tukey HSD t tests to correct for multiple comparisons.

### Distribution of LSX neurons

To quantify the distribution of neurons along the A-P, D-V, or M-L axis (Figure 1F-H), we binned the data along its range into 130μm, 220 μm, and 100 μm sized bins respectively and used the Catmull-Rom spline function implemented by Microsoft Excel to interpolate the curves of the means and SEMs. Median lines were calculated by averaging the median A-P, D-V, or M-L values of the animals in each projection population. Two-way ANOVAs with projection population and bin number as factors were used with post-hoc Tukey HSD t tests to correct for multiple comparisons when a significant interaction or main effect was found.

### Density contour plots

To create the density contour plots in Figure 2, we selected an anterior (0.845 mm), intermediate (0.545 mm), and posterior (0.02 mm) A-P coordinate at which to quantify the density and spread of the LS projection populations. When there was not a section registered at one of those exact A-P values, we chose the closest section to that A-P value. First, we made a mesh grid of 50 values with jumps of 0.02 on the x axis spanning 0 to 1 and jumps of 0.04 on the y axis spanning 0.2 to −2. These ranges were selected based on the size of one hemisphere of the LS. We then concatenated the M-L and D-V values across animals within a projection, then calculated density using the kernel smoothing *ksdensity* function in Matlab with a bandwidth of 0.08. The density values were normalized to the peak, and the normalized contours were plotted for density values ranging from 0.3 to 0.9. The centroid was quantified as the center position of the 90 percent contour values, and the spread plotted in Figure 2B,D,F was defined as the maximum height and width of the 90 percent contour. Kolmogorov-Smirnov tests were used for pairwise comparisons between the 90 percent contour of each projection population at each of the three A-P values with Bonferroni correction for multiple comparisons.

### LS projection populations by LSX subregion

To assess the distribution of neurons across the lateral septal complex (LSX) (Supplementary Figure 3C), we used the output from the Wholebrain package for each animal and quantified the number of cells labeled as belonging to the septohippocampal nucleus (SH), the septofimbrial nucleus (SF), or the rostral (LSr), causal (LSc), or ventral compartments (LSv) of the LS. These counts were normalized by animal using the total number of LSX cells labeled and plotted as the mean (± SEM) with individual animals for each projection population. A two-way ANOVA with projection population and subregion (SH, SF, LSr, LSc, LSv) as factors was used with post-hoc Tukey HSD t tests to correct for multiple comparisons.

### Number of sections included

To ensure a similar number of LSX sections were collected from each animal (Supplementary Figure 3B), we binned the data along the A-P axis into 200 μm bins and counted the number of sections within each bin, regardless of whether cell labeling was present. We then used the Catmull-Rom spline function implemented by Microsoft Excel to interpolate the curve of the means and SEMs. A two-way ANOVA with projection population and bin number as factors was used with post-hoc Tukey HSD t tests to correct for multiple comparisons due to the significant interaction and main effect found.

#### Quantification of brainwide inputs

Analysis of the whole brain inputs to LS projection populations was restricted to cells labeled in the hemisphere unilateral to the injections, ranging from 2.545 mm to −4.955 mm on the A-P axis. This measure was adopted to rule out our inability to disentangle labeling occurring from viral spread to the contralateral hemisphere for midline targets (e.g. VTA) relative to true contralateral labeling.

### Distribution of whole brain inputs

To quantify the distribution of neurons along the A-P axis (Figure 3D), we binned the data along its range (2.545 mm to −4.955 mm) into 500 μm bins and used the Catmull-Rom spline function implemented by Microsoft Excel to interpolate the curves of the means and SEMs. A two-way ANOVA with projection population and bin number as factors was used with post-hoc Tukey HSD t tests to correct for multiple comparisons when a significant main effect was found.

### Spider diagrams

To create the spider diagrams (Figure 3E), we used the output from the Wholebrain package to quantify the number of input cells across the whole brain. We first quantified the number of inputs belonging to each of the major regions as defined by the Allen Brain Atlas, specifically the hippocampal formation (HPF), isocortex (CTX), striatum (STR), pallidum (PAL), hypothalamus (HY), thalamus (TH), and midbrain (MB). Inputs were normalized by animal using the total number of input neurons counted then averaged across projection populations. The top subregions of the major regions were identified and included in the spider diagram if the number of inputs comprised at least 2% of the total inputs to that projection population. Abbreviations correspond to regions listed in the Allen Brain Atlas. A two-way ANOVA with projection population and major region (HPF, CTX, STR, PAL, HY, TH, MB) as factors was used with post-hoc Tukey HSD t tests to correct for multiple comparisons.

### Subregion barplots

To quantify the inputs from specific subregions (Figure 4, Figure 5B, Supplementary Figure 5A), cells labeled as belonging to each subregion were counted and normalized by the total number of inputs per animal. Percentages of inputs from individual animals were plotted with the mean (± SEM) for each projection population. Two-way ANOVAs with projection population and subregion as factors were used with Tukey HSD post-hoc t tests.

To analyze the percentage of inputs from dorsal and ventral CA1 and CA3 (Figure 5C-D), a cutoff of −3 mm for the D-V coordinate was implemented. If the D-V coordinate for a neuron was at or greater than −3 mm, the cell was counted as a dorsal cell, and if the D-V coordinate was less than −3 mm, the cell was counted as a ventral cell. The counts were normalized by the total number of inputs in each animal then plotted as the mean (± SEM) with individual counts. Two-way ANOVAs with projection population and depth (dorsal or ventral) as factors were used with post-hoc Tukey HSD t tests to correct for multiple comparisons.

### Hippocampal ordering by dorsoventral (D-V) coordinate

To rank the LS projection populations in Figure 5E (left panel), we organized the LS projection populations by their average median D-V coordinate from the retrograde tracing dataset and the median D-V coordinate of the CA1 inputs and the CA3 inputs from the modified rabies tracing dataset. These median D-V coordinates from the CA1 inputs and CA3 inputs were then plotted in the right panel with a line connecting the median values for the CA1 and CA3 inputs.

### Hippocampal inputs density distribution

To visualize the density of the hippocampal inputs (left panels of Figure 5F-G and Supplementary Figure 5B), we generated scatter plots of the CA1 (Figure 5F), CA3 (Figure 5G), and CA2 (Supplementary Figure 5B) where each point is colored by the spatial density of nearby points. A MATLAB function *scatter_kde* uses the kernel smoothing function to compute each point’s probability density estimate (PDE). It then uses the PDE to color each point. The + symbol represents the median M-L and median D-V coordinates from concatenated data from the three animals per projection population. We also generated normalized density plots (right panels of Figure 5F-G and Supplementary Figure 5B) of the distribution of the neurons along the dorsoventral axis (−1 mm to −6.5 mm) in 500 μm bins. The distributions for individual animals are overlaid in gray, while the mean distribution (± SEM) is in the color corresponding to the LS projection population. The gray dashed line spanning the left and right panels corresponds to the peak of the normalized density distribution plot.

### Thalamic inputs density distribution

To determine the density distribution of input cells across the thalamus (Figure 6A), we first identified all input neurons for each animal that were identified to be part of the thalamus in accordance with the Allen Brain Atlas. Next, the inputs were collapsed across three A-P ranges. The A-P range for the anterior thalamus (Figure 6A, top row) was −0.48 mm to −0.9 mm. The A-P range for the intermediate thalamus (Figure 6A, middle row) was −0.9 mm to −1.455 mm. The A-P range for the posterior thalamus (Figure 6A, bottom row) was −1.455 mm to −2.155 mm. We next binned the thalamus in 75 μm^2^ bins (M-L range 0 to 3 mm and D-V range −2 to −5.5 mm) and determined the percentage of the total inputs in each bin. We then averaged these percentages across all three animals per projection population and plotted the pixels with the color corresponding to the average percentage.

### Rank ordering of thalamic and hypothalamic inputs

To visualize the rank orders of the thalamic (Figure 6B) and hypothalamic (Figure 7A) input regions, we first identified the top ten thalamic and hypothalamic inputs across all six projection populations by rank ordering the average percentage of inputs for all thalamic and hypothalamic regions for each projection population. 14 thalamic and 15 hypothalamic nuclei were identified that constitute the top 10 regions across all projection populations. Cells that were determined to be part of the thalamus and the hypothalamus (labeled TH or HYP) but not sorted as belonging to a particular nucleus by Wholebrain were excluded from this analysis. For the thalamus, we determined what fraction of the top 14 thalamic region inputs each region represented and visualized it using treemaps (Figure 6B). For the hypothalamus, we determined what fraction of the top 15 hypothalamic region inputs each region represented and visualized it using treemaps (Figure 7A).

### Scatter plots and counts of thalamic and hypothalamic input

The scatter plots in Figures 6C-H and 7B-F represent specific regions shown in specific colors (e.g. the lateral hypothalamic neurons are shown in red in Figure 7B). The gray dots represent all thalamic (Figure 6) or hypothalamic inputs cells (Figure 7). For each plot, the data points are collapsed across all A-P coordinates and across all animals for each projection population. To create the bar plots in Figures 6C-H and 7B-F (right column), the number of neurons labeled as belonging to a specific thalamic or hypothalamic nucleus were counted for each animal and averaged across the projection population. The mean (± SEM) number of neurons for each subregion was then plotted with individual values overlaid.

#### Data visualization

To create example coronal sections of labeling in the LS (Figure 1E) or across the whole brain (Supplementary Figure 4), we used the *schematic.plot* function in the Wholebrain package to plot the cells labeled during segmentation onto coronal sections at defined A-P coordinates. Example sections within a projection population are from a single representative animal. To create the glass brain diagram (Figure 3C), we used the *glassbrain* function in the Wholebrain package to plot all inputs labeled in a representative animal. Each dot represents a single neuron and the colors correspond to those in the Allen Brain Atlas.

## Supplementary materials

**Supplementary Table 1. Statistical analyses associated with each figure.**

**Supplementary Figure 1.**
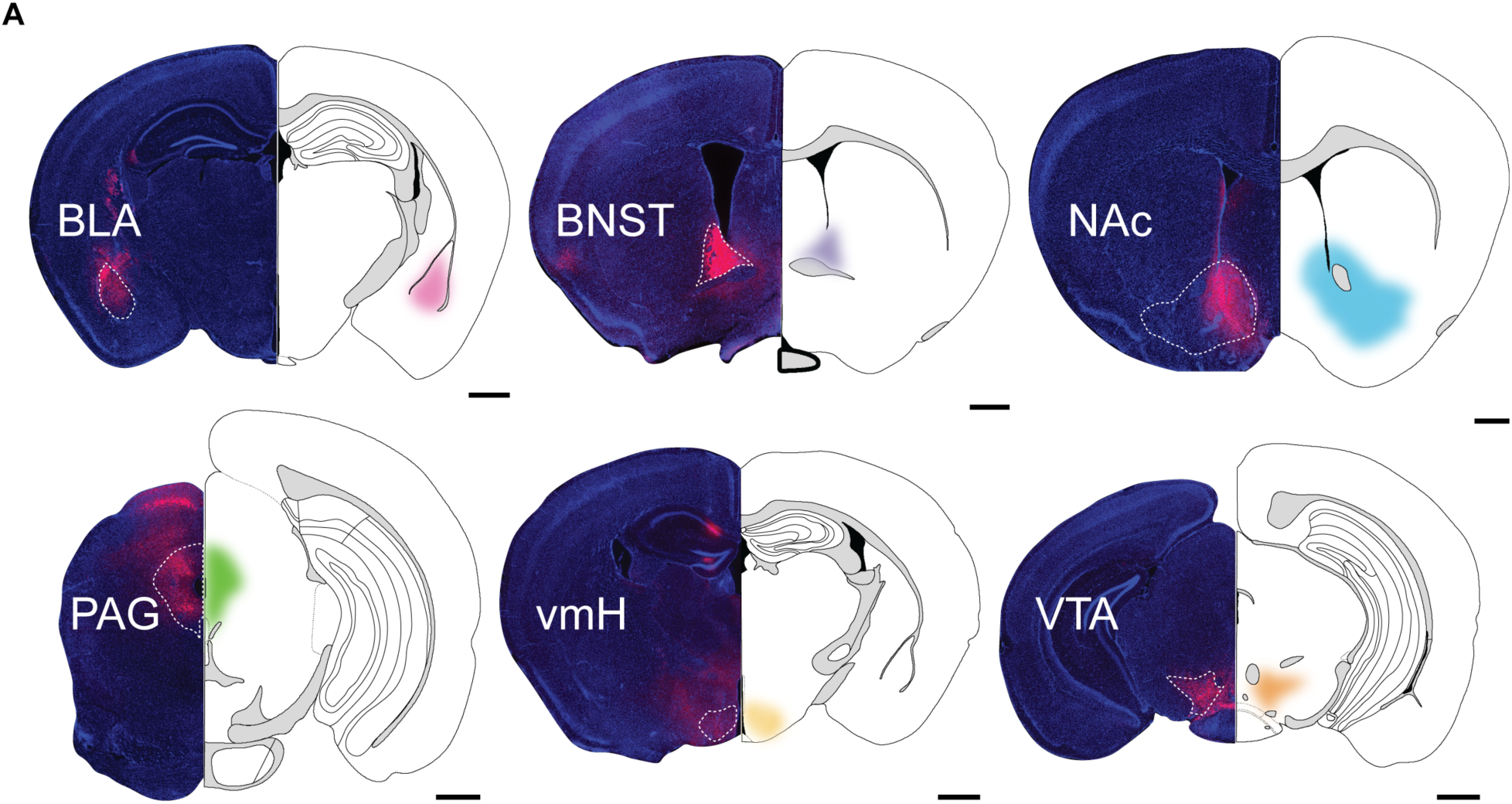
Downstream brain regions of select LS projection populations. A) Coronal sections from representative animals showing expression of the retrograde Cre virus with an mCherry tag in downstream target regions (left half). DAPI staining in blue. The corresponding atlas section is traced with the target region shaded (right half). Scale bars represent 1 mm.

**Supplementary Figure 2.**
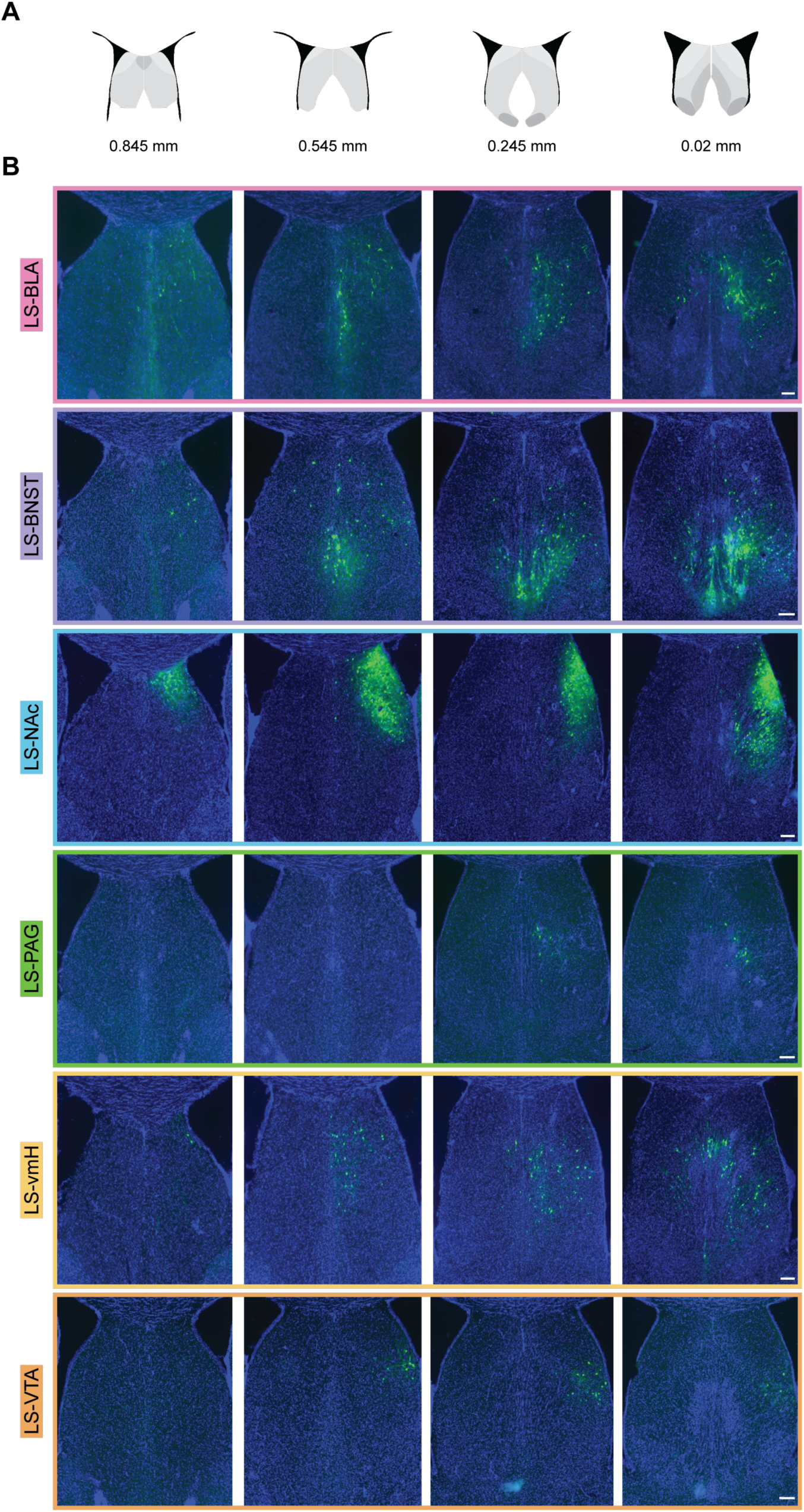
Retrogradely labeled LS projection neurons across its A-P axis. A) Schematic of coronal sections at four different A-P coordinates spanning the LS. B) Example coronal sections from representative animals showing retrogradely labeled neurons in green (GFP) that comprise the LS projection populations across four different A-P coordinates. DAPI labeling is seen in blue. Scale bars represent 100 μm.

**Supplementary Figure 3.**
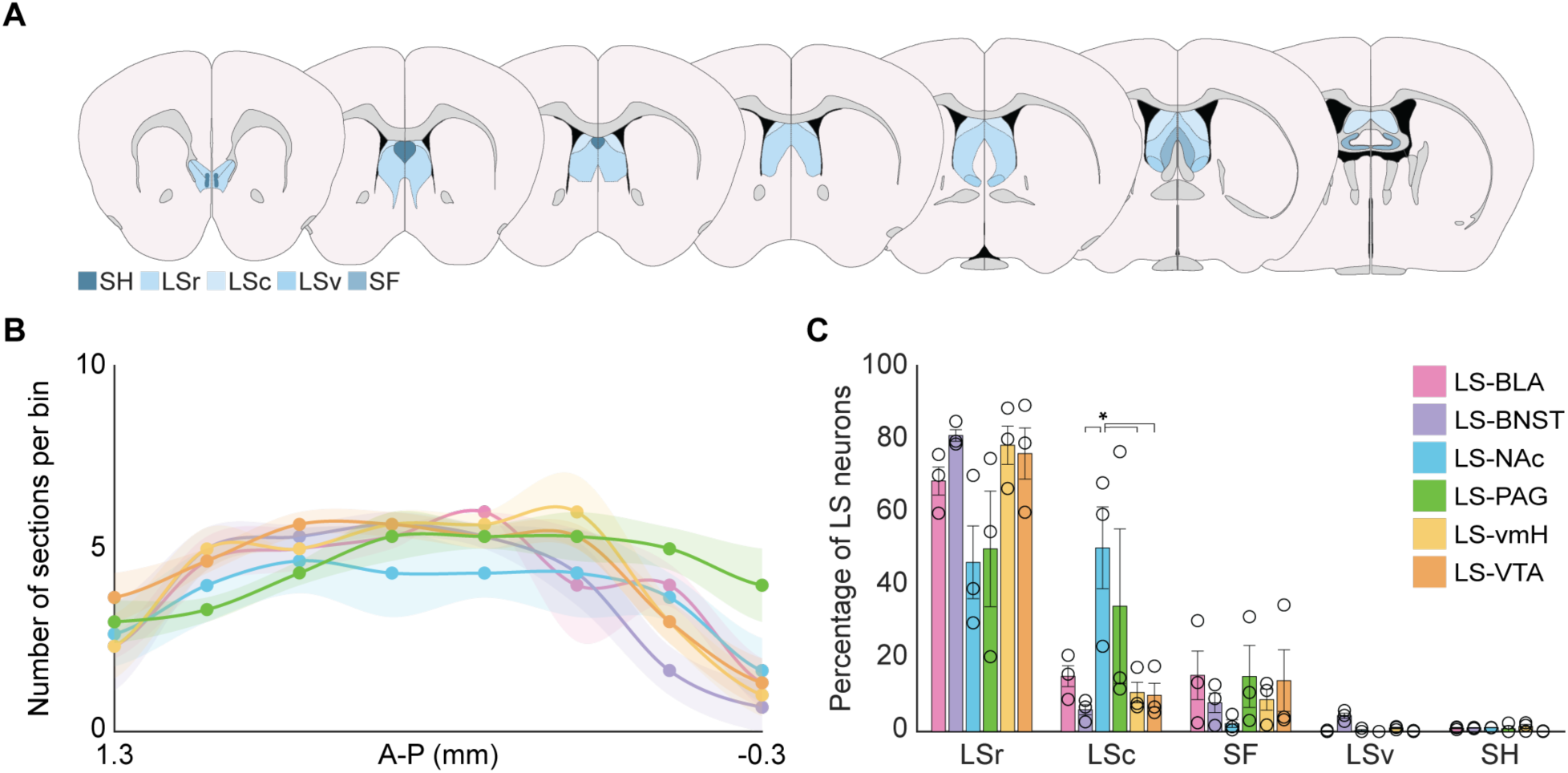
Retrogradely labeled LS projection neurons occupy different subregions within the LS. A) Schematic of coronal sections with the components of the lateral septal complex (LSX) highlighted in different colors (caudal LS (LSc), septohippocampal nucleus (SH), rostral LS (LSr), ventral LS (LSv), septofimbrial nucleus (SF)). B) Average number of sections used for cell counting across each projection population. Two-way ANOVA with projection population and bin as factors (interaction: p = 0.001, projection population: p = 1, bin: p = 3.73*10^-23^) with post-hoc t tests (see Supplementary Table 1). C) The LSr sends significantly more inputs to the BLA, BNST, vmH, and VTA populations, while the LS-NAc projection consists of more LSc neurons than the LS-BNST, LS-vmH, or LS-VTA projection populations. Two-way ANOVA with projection population and LSX subregion as factors (interaction: p = 0.001, projection population: p = 1, LSX subregion: p = 9.97*10^-25^) with post-hoc t tests comparing subregion and projection population (see Supplementary Table 1). Values from individual animals indicated by unfilled circles. Error bars and shaded error regions indicate ± SEM. N = 3 animals per projection population.

**Supplementary Figure 4.**
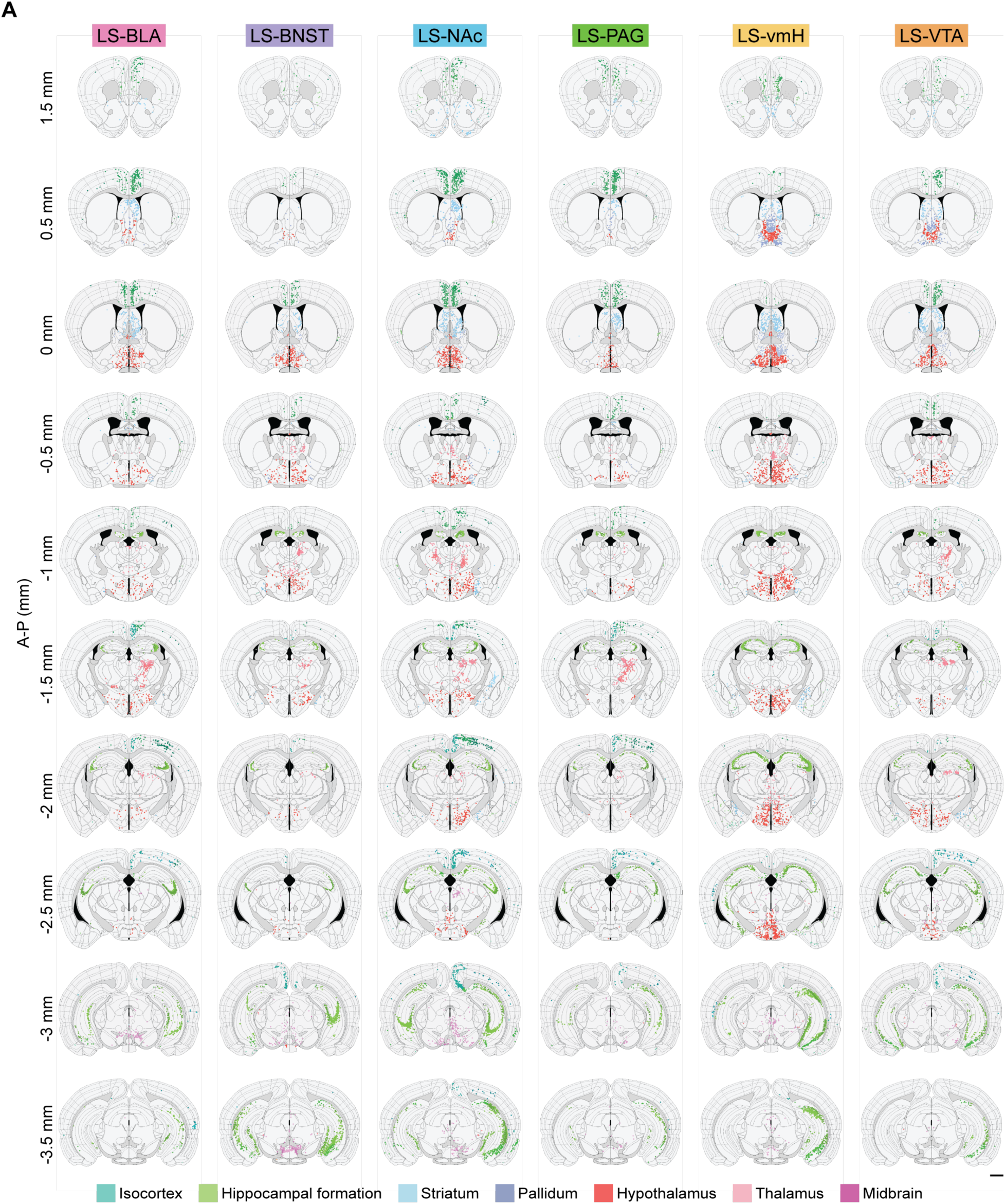
Brainwide monosynaptic inputs to individual LS projection populations in representative animals. A) Coronal sections showing registration and neuron segmentation along the A-P axis of the brain from 1.5 mm anterior to bregma to 3.5 mm posterior to bregma in representative animals. Each column corresponds to one animal from an individual projection population. Rows correspond to specific A-P coordinates relative to bregma. Each dot represents one neuron. Dot colors correspond to different input regions as defined by the Allen Brain Atlas. Scale bar represents 1 mm.

**Supplementary Figure 5.**
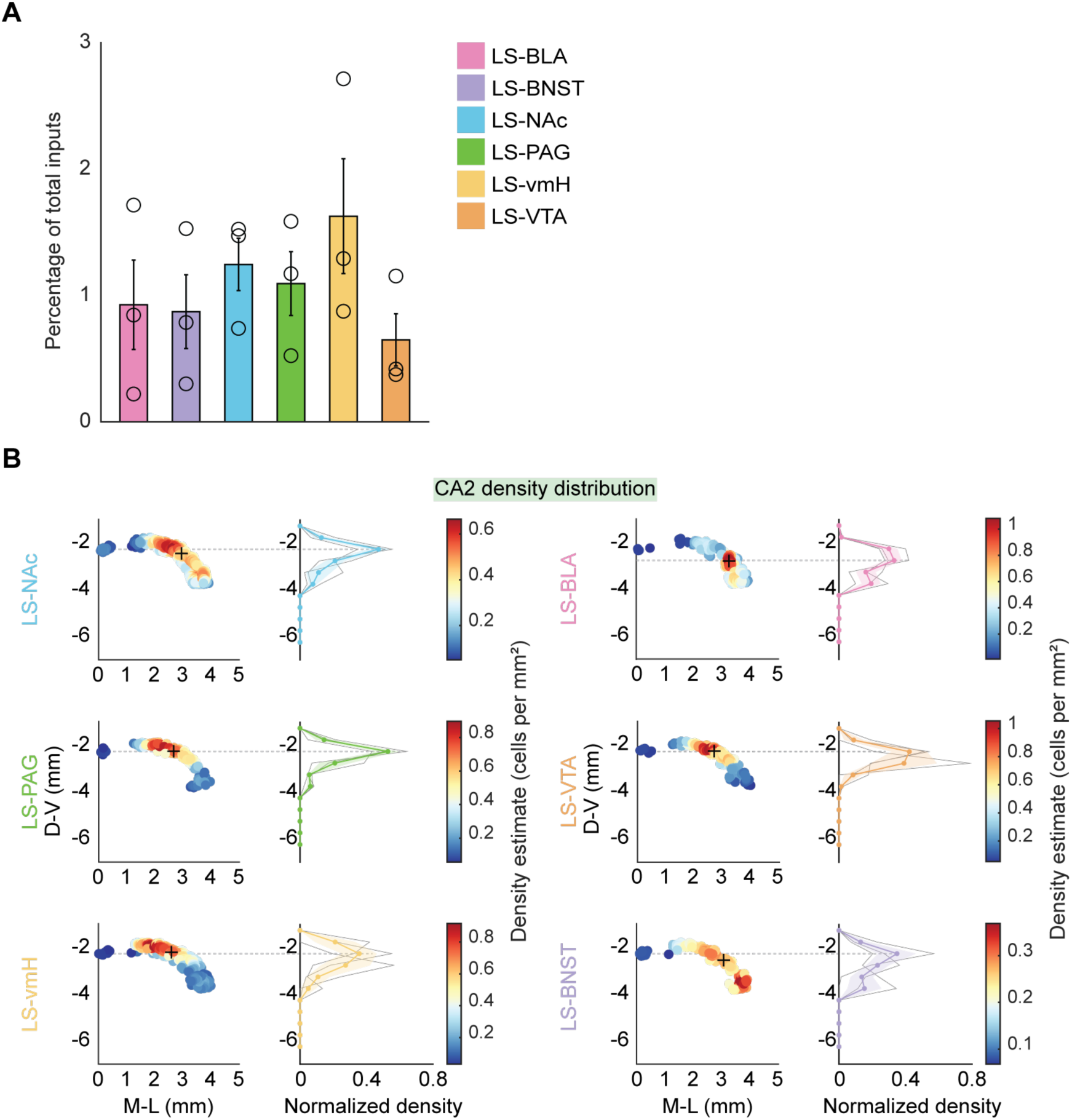
CA2 inputs to LS projection populations. A) Percentage of CA2 inputs to each projection population. B) Density plots (left column) and distribution of density (right column) of CA2 inputs along the D-V axis of the hippocampus (A-P coordinate: −1.255 mm to −3.08 mm) across all animals in each projection population. The + indicates the median M-L and D-V coordinate for the CA2 inputs to each LS projection population (+ M-L/D-V coordinates - LS-NAc: 2.95/−2.47, LS-PAG: 2.67/−2.28, LS-vmH: 2.59/−2.23, LS-BLA: 3.26/−2.82, LS-VTA: 2.70/−2.26, LS-BNST: 3.07/−2.57). The dashed gray line spanning the left and right panels corresponds to the peak of the normalized density distribution plot (D-V of gray line -LS-NAc: −2.25 mm, LS-PAG: −2.25 mm, LS-vmH: −2.25 mm, LS-BLA: −2.75 mm, LS-VTA: −2.25 mm, LS-BNST: −2.25 mm). Values from individual animals indicated by unfilled circles or gray lines. Error bars and shaded error regions indicate ± SEM. *p < 0.05. N = 3 animals per projection population.

